# 4D bioprinting shape-morphing tissues in granular support hydrogels: Sculpting structure and guiding maturation

**DOI:** 10.1101/2024.08.09.606830

**Authors:** Ankita Pramanick, Thomas Hayes, Eoin McEvoy, Abhay Pandit, Andrew C. Daly

## Abstract

During embryogenesis, organs undergo dynamic shape transformations that sculpt their final shape, composition, and function. Despite this, current organ bioprinting approaches typically employ bioinks that restrict cell-generated morphogenetic behaviours resulting in structurally static tissues. Here, we introduce a novel platform that enables the bioprinting of tissues that undergo programmable and predictable 4D shape-morphing driven by cell-generated forces. Our method utilises embedded bioprinting to deposit collagen-hyaluronic acid bioinks within yield-stress granular support hydrogels that can accommodate and regulate 4D shape-morphing through their viscoelastic properties. Importantly, we demonstrate precise control over 4D shape-morphing by modulating factors such as the initial print geometry, cell phenotype, bioink composition, and support hydrogel viscoelasticity. Further, we observed that shape-morphing actively sculpts cell and extracellular matrix alignment along the principal tissue axis through a stress-avoidance mechanism. To enable predictive design of 4D shape-morphing patterns, we developed a finite element model that accurately captures shape evolution at both the cellular and tissue levels. Finally, we show that programmed 4D shape-morphing enhances the structural and functional properties of iPSC-derived heart tissues. This ability to design, predict, and program 4D shape-morphing holds great potential for engineering organ rudiments that recapitulate morphogenetic processes to sculpt their final shape, composition, and function.

## 1. Introduction

Bioprinting technology holds tremendous potential for developing artificial tissues and organs that mimic the complexity of their native counterparts ^1–4^. Through layering of cell-laden bioinks using extrusion or lithography technology, bioprinting technology enables spatial patterning of multiple cell populations into tissue-like constructs ^5–8^. Notable advancements in embedded bioprinting, a technique where the bioink is extruded into a supportive hydrogel to prevent filament collapse under gravitational or surface tension forces, have enabled the fabrication of anatomically accurate constructs mimicking the geometry of an entire heart ^9,10^. More recently, the embedded bioprinting method has been employed for the direct extrusion of human induced pluripotent stem cell (hiPSC)-derived cardiomyocytes into cardiac tissue constructs that display organised sarcomeres and contractile capacity ^11^. While bioprinting technologies can create tissue-like constructs with precise geometries that mimic the anatomical shape of human organs, the encapsulated cells within these constructs remain structurally and functionally immature compared to their native counterparts. This limitation hinders the application of bioprinted tissues in drug screening and regenerative medicine.

Traditional bioprinting approaches typically focus on replicating the final geometrical structure of the organ, often using bioinks with high crosslinking and/or polymer network densities ^12,13^. This results in static cell-laden hydrogels that undergo limited post-printing shape changes. This restricts embedded cells from initiating and engaging in self-organising behaviours that are essential for evolution into functional tissues. This contrasts sharply with organogenesis, where cellular activities like proliferation and contractile forces drive dynamic shape-morphing patterns that are crucial for maturation ^14^. Recent advances in materials science have led to the development of 4D materials capable of undergoing pre-programmed shape-transformations when exposed to specific environmental cues like magnetic fields, light, and chemical degradation ^15–18^. As an example, composite hydrogels embedded with aligned cellulose fibers have been shown to exhibit anisotropic swelling, which can be spatially controlled to program shape-morphing in 3D printed structures ^17^. These advancements have led to the emergence of 4D bioprinting, where cell-laden hydrogels are engineered to execute pre-determined shape transformations ^19–22^. For example, cell compatible alginate and hyaluronic acid hydrogels can be engineered to undergo 4D shape-morphing triggered by spatial patterns of swelling ^23–25^. Although promising, these shape changes are artificially induced by hydrogel swelling and are not coupled or driven by cell-generated processes, as occurs in biological tissues.

Recent research has shown promise in fibrillar bioinks that can shrink and shape-morph under active cell-generated forces within soft support hydrogels ^26,27^. For example, collagen bioinks extruded into simple beam structures display structural instabilities and shape-morph as cells contract the collagen network ^28^. In another example, fibrillar bioinks composed of electrospun fibers were shown to shrink as cells contracted the fibers, with spatial variations in fiber density driving patterns of contractile behavior ^27^. However, translating and scaling these promising 4D approaches to organ bioprinting faces significant challenges. A primary challenge lies in the limited understanding of the physical principles that govern 4D shape-morphing in bioprinted constructs. A fundamental understanding of these physical principles is essential for the design and optimisation of hydrogel platforms that can predictably program and modulate 4D shape-morphing across the length scales relevant to organ development. For example, key knowledge gaps include the impact of bioink and support bath stiffness or viscoelasticity on shape-morphing. We also lack a clear understanding of how the shape and geometrical features of the initial bioprinted construct influence the development of mechanical instabilities and subsequent shape changes. Additionally, the influence of ECM composition and cell phenotype on shape-morphing remains poorly understood. Crucially, the impact of 4D shape-morphing on cell morphology, differentiation, and maturation has not been explored. These knowledge gaps are key barriers preventing the design of materials that can effectively guide and modulate 4D shape-morphing, as well as the creation of bioprinted constructs capable of undergoing programmed morphogenetic behaviors essential for their structural and functional evolution into mature tissues.

Here, we developed a bioprinting platform that enables the fabrication of tissues that undergo predictive and programmable 4D shape-morphing via cell-generated forces. Utilizing embedded bioprinting, we deposit collagen-hyaluronic acid bioinks into yield-stress granular support hydrogels, which can locally fluidize to accommodate and regulate tissue shape transformations. We systematically investigated key variables influencing the shape-morphing process, including initial print geometry, bioink ECM composition, cell phenotype, and support hydrogel viscoelasticity. Importantly, we also developed a computational finite-element model that accurately predicts tissue shape-morphing patterns and internal changes in cell and ECM organisation during morphing. As a proof-of-concept, we demonstrate that programmed 4D shape-morphing enhances the structural and functional properties of bioprinted iPSC-derived heart tissues. This platform holds significant promise for bioprinting organ rudiments that undergo programmed shape changes to sculpt their final composition, form, and function.

## 2. Main

### 2.1. 4D bioprinting shape-morphing tissues in granular support hydrogels

To establish our 4D bioprinting platform, we first developed an ECM bioink capable of supporting shape-morphing via cell-generated forces. Collagen type I was chosen for its ability to facilitate cell attachment and contraction, and hyaluronic acid was added to enhance viscosity and impart shear-thinning properties necessary for extrusion bioprinting (Figure 1a, Figure S1a). Next, we used embedded bioprinting to deposit the ECM bioink into granular support hydrogels composed of agarose microgels (Figure 1a i). The shear-thinning and self-healing properties of this support hydrogel enabled high-resolution printing of the ECM bioink into geometries such as tubes or spirals (Figure 1a ii, Movie S1, S2, S3). The addition of hyaluronic acid improved the resolution of printed filaments within the support bath (Figure S1a). For initial studies, human cardiac fibroblasts (CFs) were encapsulated within the bioink. Following deposition into the support hydrogels, bioprinted constructs underwent significant shape morphogenesis over 14 days of culture (Figure 1a ii). Notably, the initial print geometry strongly influenced the shape-morphing patterns, with bioprinted tubes exhibiting isotropic shrinkage and bioprinted spirals gradually unfolding (Figure 1a ii, Figure S1b ii, iii). Acellular prints without CFs did not exhibit shape-morphing, confirming the cell-mediated nature of the process (Figure S1c). Importantly, cell viability remained high within the morphing tissues, and nanoindentation revealed a tenfold increase in elastic modulus over the culture period (Figure S2a). This is likely due to increased cell density and ECM compaction, which was evident in confocal images (Figure 1a). Immunofluorescence staining for fibronectin also revealed that encapsulated cells synthesized nascent ECM proteins during culture, which will further contribute to the evolving tissue mechanics (Figure S2b).

**Figure 1.**
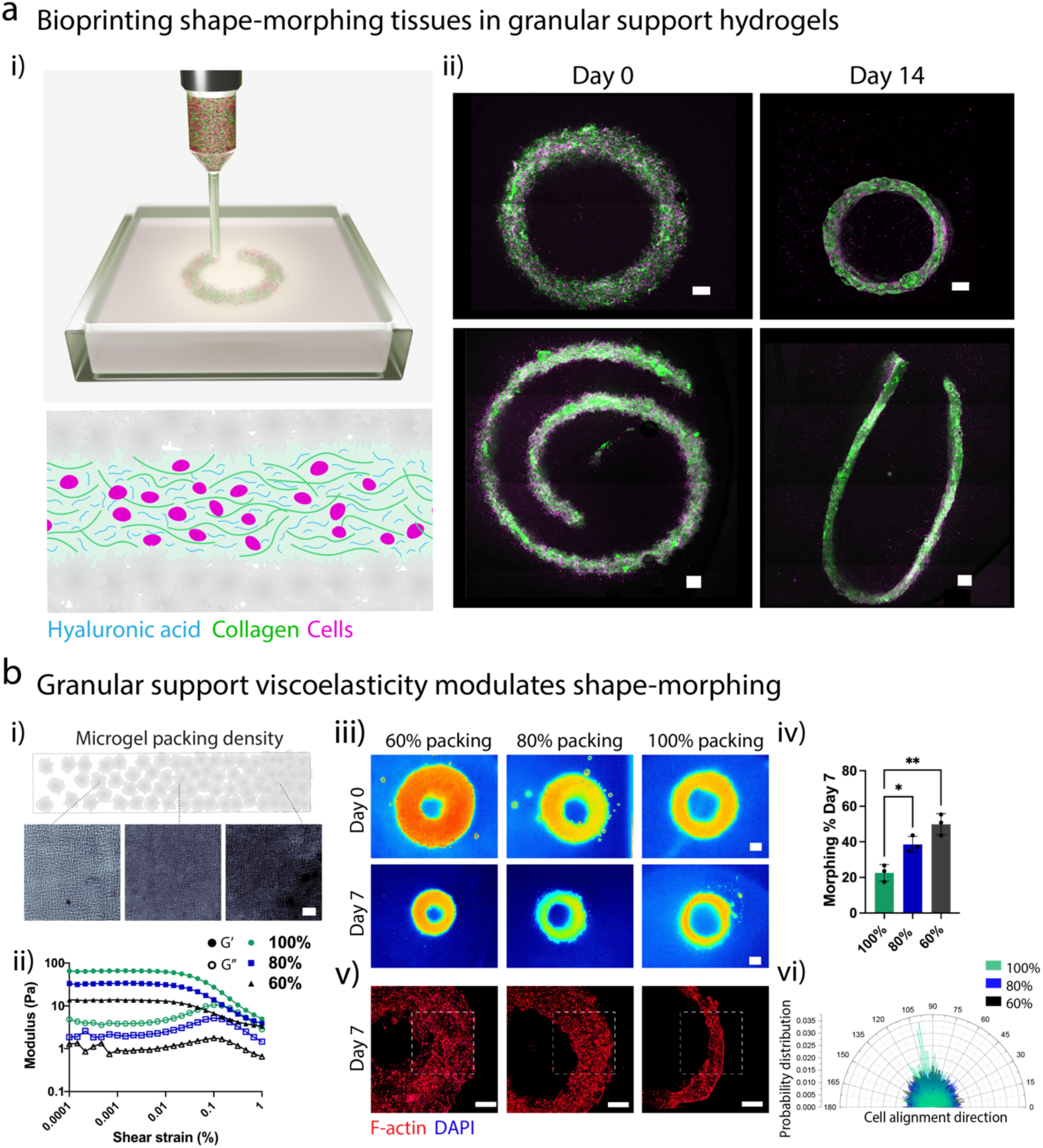
Embedded bioprinting of shape-morphing tissues within granular support hydrogels: **(a)** (i) Schematic representation of embedded bioprinting platform, and (ii) stitched confocal images of bioprinted tubes and spirals cultured in the support hydrogel over 14 days of culture. Scale bars 500µm. Images are representative of n=3 biological replicates. **(b)** (i) Schematic representation and brightfield images of granular support hydrogels with varied packing density (Scale bars 500µm), and (ii) rheological characterisation of storage and loss modulus at varying shear strain (0.0001-1%) as a function of support hydrogel packing density. (iii) Coloured brightfield images demonstrating tissue shape-morphing in granular support hydrogels with varied viscoelasticity (60, 80%, 100% packing density) at day 0 and day 7. Scale bar 1mm. (iv) Quantification of shape-morphing on day 7 in different granular support hydrogel formulations based on brightfield images (n=3 biological replicates, one-way ANOVA with Tukey’s multiple comparison test, where * denotes p < 0.05, and ** denotes p < 0.01). (v) Confocal images of actin-stained cells within shape-morphing tubes cultured in support hydrogel formulations with varied viscoelasticity, scale bar 500µm, and (vi) polar histograms of cell alignment within the morphing tissues (n=3 biological replicates per formulation).

Next, we investigated the impact of support hydrogel rheology on tissue shape-morphing. By adjusting microgel packing density through serial dilutions (from 100% to 60%), we modulated the storage modulus of the support hydrogel from approximately 80 Pa to 10 Pa, while maintaining its shear-thinning and self-healing properties (Figure 1b i, ii, Figure S2c i). Notably, softer support hydrogel formulations resulted in significantly greater shrinkage of the bioprinted tubes over 7 days of culture (Figure 1b iii, iv). Interestingly, support hydrogel viscoelasticity also impacted internal cell organisation within the morphing tissues (Figure 1b v). In softer support hydrogels, printed cells were circular and randomly orientated, whereas, in stiffer supports, we observed increased cell spreading and alignment along the principal tissue axis (Figure 1b v, vi). This demonstrates that support hydrogel rheology can actively modulate shape morphogenesis at both the cell and tissue scales. We hypothesize that higher packing density formulations offer greater resistance to shrinking tubes, leading to higher internal tissue stresses and strains experienced by cells, thereby promoting cell alignment. This phenomenon aligns with the use of physical constraints like micropillars to guide alignment in collagen microtissues ^29–31^. Our results highlight the potential of granular support hydrogels to act as dynamic constraints for bioprinted tissues with complex 3D geometries, offering a means to provide mechanical stimuli to larger-scale constructs that cannot be effectively actuated using conventional micropillar techniques. This expands the utility for support hydrogels beyond merely preventing bioink collapse during printing, and demonstrates that their properties can be modulated to guide tissue maturation post-bioprinting. Having established a platform for 4D bioprinting shape-morphing tissues in granular support hydrogels, we next focused on developing a computational model to predict and design tissue shape evolution.

### 2.2. Finite element modelling to predict 4D shape-morphing of bioprinted tissues

To gain a deeper understanding of the mechanical behaviour of bioprinted constructs during 4D shape-morphing, we next focused on developing a finite element model capable of predicting tissue shape evolution. To do this, we implemented an active modelling framework that captures cell active contractility, remodelling, and alignment using a thermodynamically consistent model for stress-fibre remodelling introduced by Reynolds et al. (Figure 2a) ^32^. Cells are embedded in a 3D anisotropic collagen gel, modelled using a hyperelastic framework where an initially uniform distribution of fibres can realign in response to active contractility. Full details of the framework are provided in the methods and supplementary section. We proceeded to investigate the shape-morphing behaviour of classic geometries (Figure 2). Simulation of a bioprinted tube to day 14 steady state led to significant construct contraction, with a 45.7% reduction of the outer diameter, in agreement with experimental findings (Figure 2b i-ii, Figure S3, Movie S4). These predictions were next validated through comparison with additional printed shapes (square, triangle, junction) (Figure 2b i-ii, Figure S3, Movie S5, S6, S7). Notably, the circularity of the internal boundary increases due to confinement imposed by the support bath hydrogel, in further agreement with observed behaviour (Figure 2b i-ii). This leads to a high stretch at the tissue vertices and, thus, a locally increased active stress. Interestingly, our model also captures the widening of a bioprinted L-shaped junction during contraction, which primarily arises due to pressure generated during compression of the support bath hydrogel (Figure 2b ii). Our analysis suggests that a reduction in granular hydrogel stiffness (analogous to packing density in Figure 1b) leads to an increase in tissue contraction and shape-morphing (Figure 2b iii-iv). Furthermore, our models predict higher internal tissue active stresses in tissues undergoing shape-morphing within stiffer support bath formulations (Figure 2b iii). This result helps explain the increased cell spreading and alignment observed in higher packing density support bath formulations (Figure 1b v), as mechanical constraints have been previously shown to guide cell alignment in collagen matrices ^32^.

**Figure 2.**
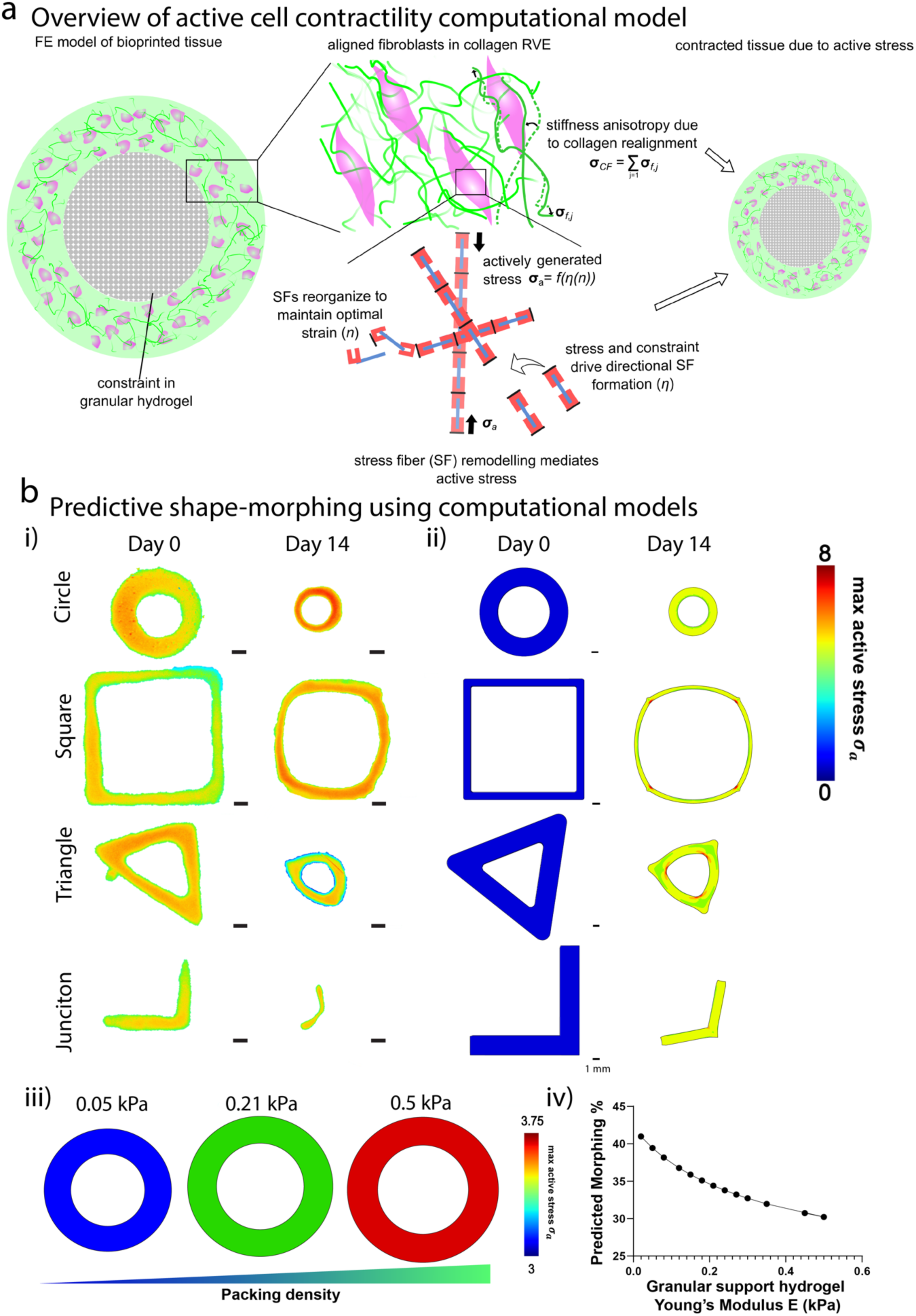
Finite element model for predicting tissue shape-morphing based on cell contractility: **(a)** Overview of finite element model that predicts shape-morphing using thermodynamically-motivated reorganisation of cells and stress fibres to maintain optimal strain within collagen matrices. (1) Cells generate active stress within the collagen matrix, (2) Active stress and physical constraints from the collagen matrix drive directional stress fibre formation, (3) Remodelling of stress fibres regulates active cell stress, and (4) Stress fibres then reorganise to maintain optimal strain. (b) Validation of FE model for predicting 4D shape-morphing of different tissue geometries (tube, square, triangle and junction) including (i) Brightfield images of experimental outcomes on day 0 compared to day 14 (scale bar 1mm), (ii) FE models of tissue shape-morphing with active stress (*σ_a_*) contour plots. (iii) Predictions for tissue shape-morphing and active stress when cultured with support baths with increasing packing density (*E* = 0.05 *kPa*, 0.21 *kPa*, 0.5 *kPa*), and (iv) Simulation predictions for tissue shape-morphing in granular support hydrogels with increasing Young’s modulus. Experimental images are representative of n=3 biological replicates.

Our computational finite element models, incorporating active cell contractility, effectively predict 4D tissue shape-morphing, providing a framework for the predictive design of shape evolution in bioprinted tissues for the first time. This is significant given the established role of active cell contractility in driving morphogenetic tissue flows during organ development ^14,33^. Our bioprinting platform, coupled with this predictive FE modelling framework, offers a powerful tool for designing and fabricating tissues that undergo programmable shape-morphing to sculpt their final form. Building upon this foundation, we next investigated how cell phenotype and ECM composition could modulate tissue shape-morphing.

### 2.3 Influence of cell phenotype and ECM composition on 4D shape-morphing

Given that 4D shape-morphing behaviours were driven by cell-generated contraction forces, we next explored how cell density, phenotype, and ECM concentration could influence the magnitude of shape change. Using a full-factorial design of experiments approach, we examined the impact of cell density and collagen concentration on the shape-morphing of CF-laden bioprinted tubes (Figure 3a i). Increased tube shrinkage (up to 42%) was observed with higher cell densities and lower collagen concentrations (Figure 3a i). FE simulations, varying cell density and bioink modulus (analogous to collagen concentration), corroborated these experimental observations, predicting greater shape-morphing with higher cell densities or lower bioink moduli (Figure 3a ii). These results demonstrate the tunability of tissue shape-morphing through controlled bioink cell and ECM composition.

**Figure 3.**
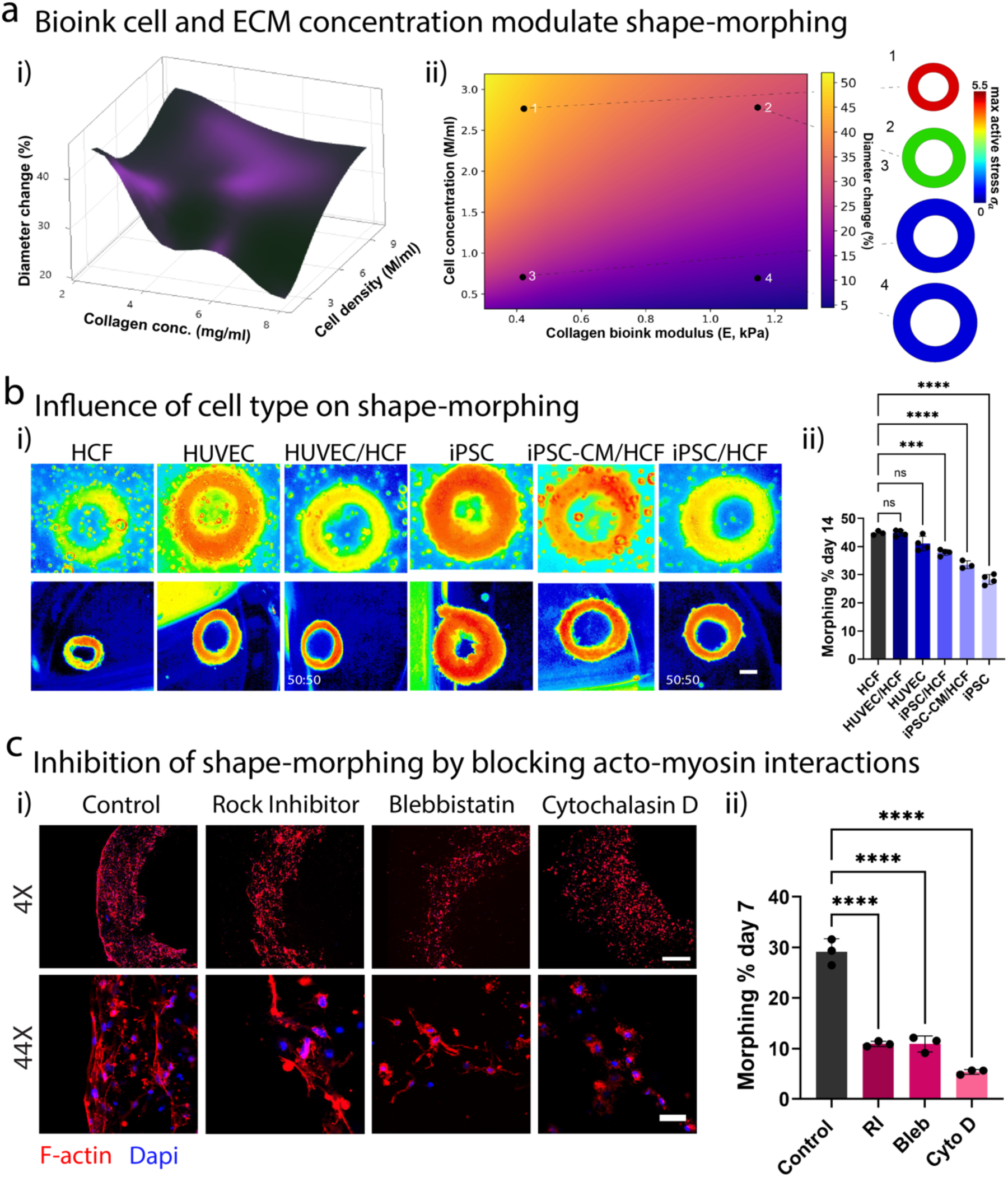
Modulating 4D shape-morphing via bioink composition and cell phenotype: **(a)** (i) Influence of bioink collagen concentration (2.4-8mg/ml) and cell density (2-10M/ml) on tissue shape-morphing demonstrated by surface plot of tube shape-morphing (% diameter change) based on full-factorial design of experiments. (ii) Computational predictions for tube shape-morphing (% diameter change) as a function of cell concentration and collagen bioink Young’s modulus, along with sample active stress (*σ_a_*) contour plots for different bioink formulations. **(b)** Influence of cell type on shape-morphing indicated by (i) coloured brightfield images at day 0 and day 14 (scale bar 1mm), and (ii) quantification of shape-morphing (% diameter change) on day 14 for each bioink cell composition (n=3/4 biological replicates, one-way ANOVA with Tukey’s multiple comparison test where ns: not significant, *** denotes p < 0.001, **** denotes p < 0.0001). **(c)** Influence of actomyosin contractility inhibitors (Rock Inhibitor Y27632 – 10µM, Blebbistatin – 20µM, and Cytochalasin D – 10µM) on tissue shape-morphing demonstrated by confocal images of bioprinted tubes at day 7 at low (scale bar 500µm) and high magnification (scale bar 50µm), and ii) quantitative measurement of shape-morphing (% diameter change) in control and inhibitor-treated bioprinted tubes (n=3 biological replicates, one-way ANOVA with Tukey’s multiple comparison test where ns: not significant, **** denotes p < 0.0001). All images are representative of at least n=3 biological replicates.

Next, we investigated the impact of cell phenotype on shape-morphing by bioprinting tissue tubes using cells known to display varying levels of actomyosin contractility: human umbilical cord-derived endothelial cells (HUVECs), human induced pluripotent stem cells (iPSCs), and iPSC-derived cardiomyocytes (Figure 3b, Figure S4b). Fibroblasts and HUVECs generated the greatest shape change (44% and 41% respectively), while iPSCs and iPSC-cardiomyocytes produced significantly less morphing (Figure 3b i, ii). Co-culturing fibroblasts with each cell type (1:1 ratio) partially restored the extent of shape-morphing compared to the respective monocultures (Figure 3b i, ii). Notably, the temporal dynamics of shape-morphing also varied across cell types, with HUVECs displaying rapid initial shape changes and fibroblasts exhibiting more consistent morphing over time (Figure S4b ii). For these cell-type studies, matrigel (5% volume) was incorporated into the bioink as it was required to support iPSC survival (Figure S4a i). iPSC survival further improved when co-cultured with fibroblasts (Figure S4a i). Confocal microscopy revealed that iPSCs formed embryoid body-like structures when cultured in the bioinks (non-printed conditions), with significant increases in embryoid body formation observed in fibroblast co-cultures (Figure S4a ii, iii). Most of these embryoid body structures were positive for OCT4 after 7 days of culture in the bioink, indicating maintenance of pluripotency (Figure S4a iv). However, when iPSCs were printed in the support hydrogel using this bioink and cultured for 7 days (i.e. shape-morphing conditions) OCT4 staining reduced indicating loss of pluripotency and differentiation (Figure S4b i). HUVEC viability was high when cultured in the bioinks for 7 days, and interconnected networks were observed in bioprinted shape-morphing tubes, particularly in fibroblast co-cultures (Figure S4a v, Figure S4b i). These results suggest that cell-type-specific differences in shape-morphing are primarily driven by variations in cell contractility, rather than variations in cell viability.

To further validate our findings and elucidate the cellular mechanisms underlying tissue morphing, we investigated the role of cell-generated forces by inhibiting actomyosin contractility and cell-ECM interactions. To do this, we treated cells with inhibitors prior to bioprinting, namely rock inhibitor (RI, 10µM: inhibits the phosphorylation of regulatory myosin light chain), blebbistatin (Bleb, 20µM: myosin heavy chain inhibitor), and cytochalasin D (cyto D, 10µm: depolymerization of actin). Notably, inhibitor-treated constructs exhibited significantly less shape-morphing (5-10%) compared to untreated controls (29%) (Figure 3c i, ii). Confocal microscopy revealed that the inhibitors induced changes in cell morphology, with blebbistatin treatment inducing dendritic morphologies and cytochalasin D treatment resulting in rounded morphologies with impaired actin spreading (Figure 3c i). Importantly, the inhibitor concentrations employed did not affect cell viability (Figure S5a). These results confirm that cell-generated forces, particularly actomyosin contractility, play a central role in driving the observed 4D shape-morphing behaviour in our bioprinted tissues. Furthermore, they demonstrate that shape-morphing can be modulated by tuning cell density, phenotype, and ECM composition, providing a modular framework for designing tissues that undergo predictable and programmable shape-morphing. Next, we investigated the impact of 4D tissue shape-morphing on internal cell and ECM organisation.

### 2.4 4D shape-morphing sculpts cell and ECM organisation in bioprinted tissues

Next, we investigated the influence of tissue shape-morphing on internal cell and ECM organisation by bioprinting tissues using FITC-conjugated collagen and TRITC-phalloidin labelled fibroblasts. High-resolution confocal microscopy and Orientation-J analysis were used to reconstruct and quantitatively assess cell and collagen organisation (Figure 4a i, Figure S5b). Comparing day 0 and day 14 samples revealed a dynamic evolution of organisation during the morphing process (Figure 4a i, ii, Figure S6a i, ii). Initially rounded and randomly oriented, cells gradually aligned circumferentially along the principal tissue axis as the tissue morphed under cell-generated forces (Figure 4a i, ii, Figure S6a i, ii). This alignment emerged regardless of the initial cell density, with higher densities leading to greater alignment (Figure S6b, S7a i, ii). Further, circumferential alignment was observed in bioprinted tubes printed with both concentric and rectilinear infill paths (Figure S8a). This, combined with the absence of cell and collagen alignment at day 0, indicates that the emergence of tissue structure was a direct consequence of the shape-morphing process, rather than shear-induced alignment that can occur during extrusion bioprinting under specific conditions ^34,35^. Importantly, cell and ECM alignment along the principal tissue axis also emerged in shape-morphing wave geometries, demonstrating the generalisable nature of the phenomenon (Figure 4b i, ii, vi, vii S8b). The FE model predictions for cell and ECM alignment, which are based on the thermodynamically driven reorganisation of stress fibres and collagen fiber remodelling, were in close agreement with the experimental results (Figure 4b iii, iv, viii). The model predicted that stress fibres and cells primarily align in the direction of principal strain (i.e. direction of maximum stretch) in the shape-morphing tissues (Figure 4b v, viii). These results strongly suggest that cytoskeletal reorganisation and alignment are driven by constraint and strain anisotropy, which emerges due to the influence of strain and active tension on the chemical potential of assembled stress fibers (see Methods Section for further details).

**Figure 4.**
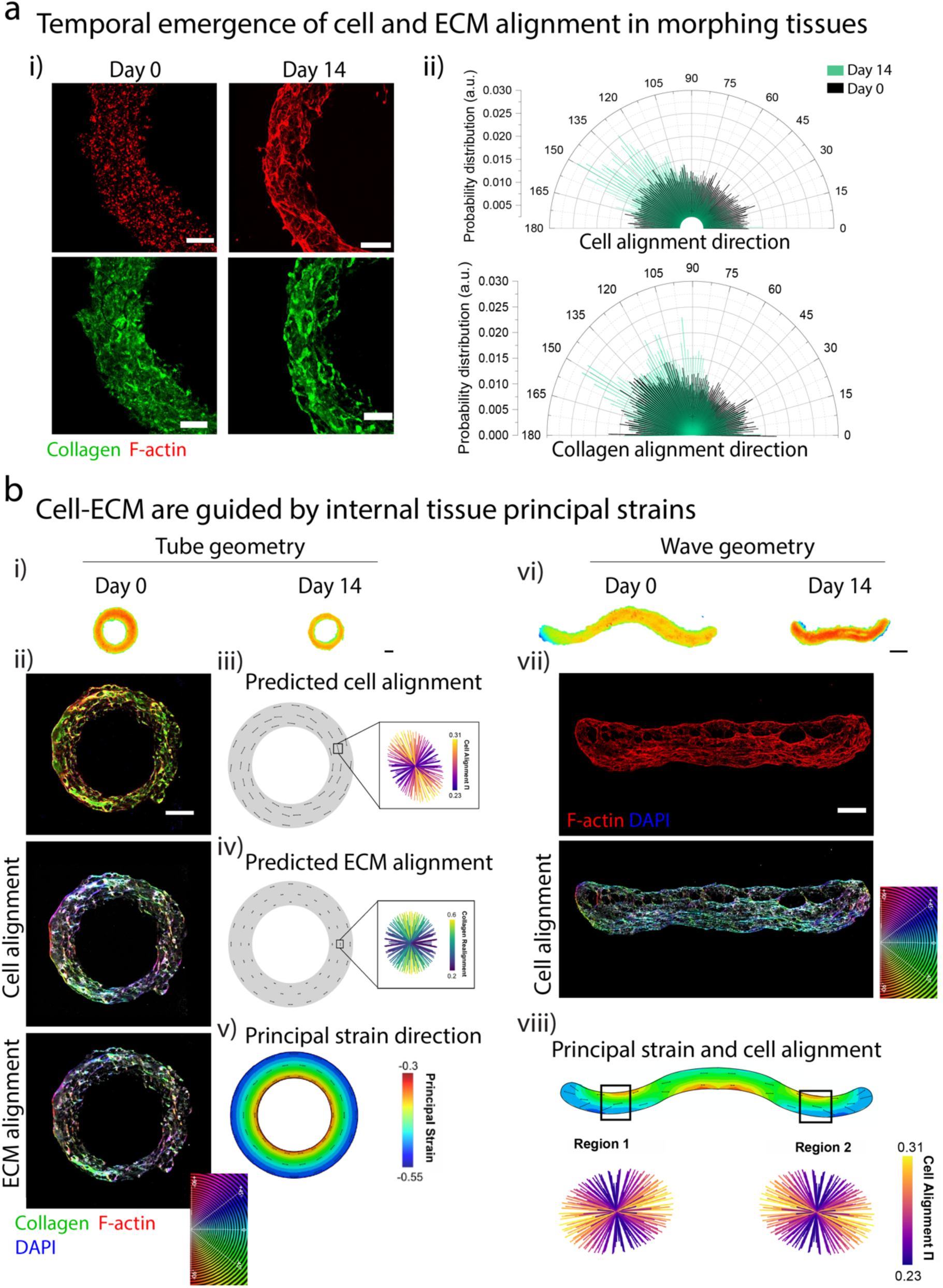
Temporal emergence of cell and ECM alignment in shape-morphing tissues: **(a)** Temporal emergence of cell and ECM alignment in shape-morphing tubes (2M/ml fibroblast cell density) over fourteen days of culture demonstrated by (i) confocal images of cells and collagen (scale bar 500µm) and (ii) polar histogram revealing the directional distribution of cells and collagen fibres with respect to the horizontal axis (n=3 biological replicates). **(b)** (i) Coloured brightfield images demonstrating shape-morphing in closed tube geometry (scale bar 1mm) along with (ii) colour mapping of cell and ECM directionality extracted from confocal images, (iii) predicted cell alignment (Π_i_ = η_i_n_i_) from the FE model, (iv) predicted ECM alignment from the FE model, and (v) contour plot of simulated principal strain magnitudes and directionality from the FE model. (vi) Coloured brightfield images demonstrating shape-morphing in open wave geometry (scale bar 1mm) along with (vii) colour mapping of cell directionality extracted from confocal images, (viii) contour plot of simulated principal strain magnitudes and directionality, merged with directionality and degree of predicted cell alignment in two highlighted regions of the tissue. All scale bars of confocal images are 500µm unless otherwise stated. Images and results are representative of n=3 biological replicates.

These results demonstrate that cell-generated forces not only drive tissue shape-morphing but also actively sculpt internal cell and ECM organisation. Importantly, we identify a general principle: alignment consistently arises along the maximum principal stress direction, likely due to stress fibre reorganisation to minimise cellular stress. Recapitulating the structural anisotropies found in mature tissues is a significant challenge in bioprinting. While shear-induced alignment has been employed to enhance cell organisation in bioprinted tissues ^34–36^, these approaches are challenging to adapt to more complex multi-axial 3D architectures. Here, we present a novel developmentally-inspired approach where bioprinted tissues are programmed to undergo shape transformations that actively sculpt their internal architecture. Since alignment is dictated by the maximum tissue strain directions that arise during morphing, this method holds great potential for recapitulating the complex anisotropies found in mature organs. Importantly, our FE modelling framework enables the prediction and design of final cell and ECM architecture a priori based on the principal strain patterns that will emerge during morphing. Building on these principles, we next explored how cell-mediated 4D shape-morphing could influence the maturation of bioprinted iPSC-derived heart tissues.

### 2.5 4D bioprinting structurally organised heart tissue models using iPSC-cardiomyocytes

Next, we leveraged our bioprinting platform to explore how cell-mediated 4D shape-morphing could influence the structural and functional maturation of iPSC-CM derived heart tissues. iPSC-CMs are structurally and functionally immature compared to adult CMs, exhibiting disorganized sarcomeres and weak, irregular contractions (Figure S9) ^37^. While recent studies have shown that iPSC-CMs can be bioprinted within collagen-based bioinks to create heart tissue models ^9,11^, the structural maturation of these tissues falls well below that of adult heart tissue. For example, in the adult heart, CMs display highly compacted and aligned morphologies optimised for electrical synchronisation and contractile output. Based on our prior results demonstrating shape-morphing guided cell alignment, combined with the importance of shape transformations in embryonic heart development ^38–40^, we hypothesized that cell-mediated 4D shape-morphing could enhance the structural maturation of bioprinted iPSC-CM derived heart tissues. We bioprinted heart tissue rings composed of a 7:3 ratio co-culture of iPSC-CMs and CFs at a high cell density of 20 million cells/ml to replicate the dense cellular nature of the myocardium. This cell number ratio was used to reflect the relative volume occupancy of CMs and CFs in the native myocardium ^41,42^.

In bioprinted tubes, the presence of CFs significantly enhanced cell-mediated tissue morphing compared to iPSC-CM monoculture controls, resulting in a fourfold increase in tube shrinkage over 28 days of culture (Figure 5a i-iii, Movie S9-11). Immunofluorescence staining for cardiac troponin T and sarcomeric alpha-actinin revealed substantial improvements in the structural organisation of iPSC-CMs within morphing tissues (Figure 5a i, ii, Figure S10a i).

**Figure 5.**
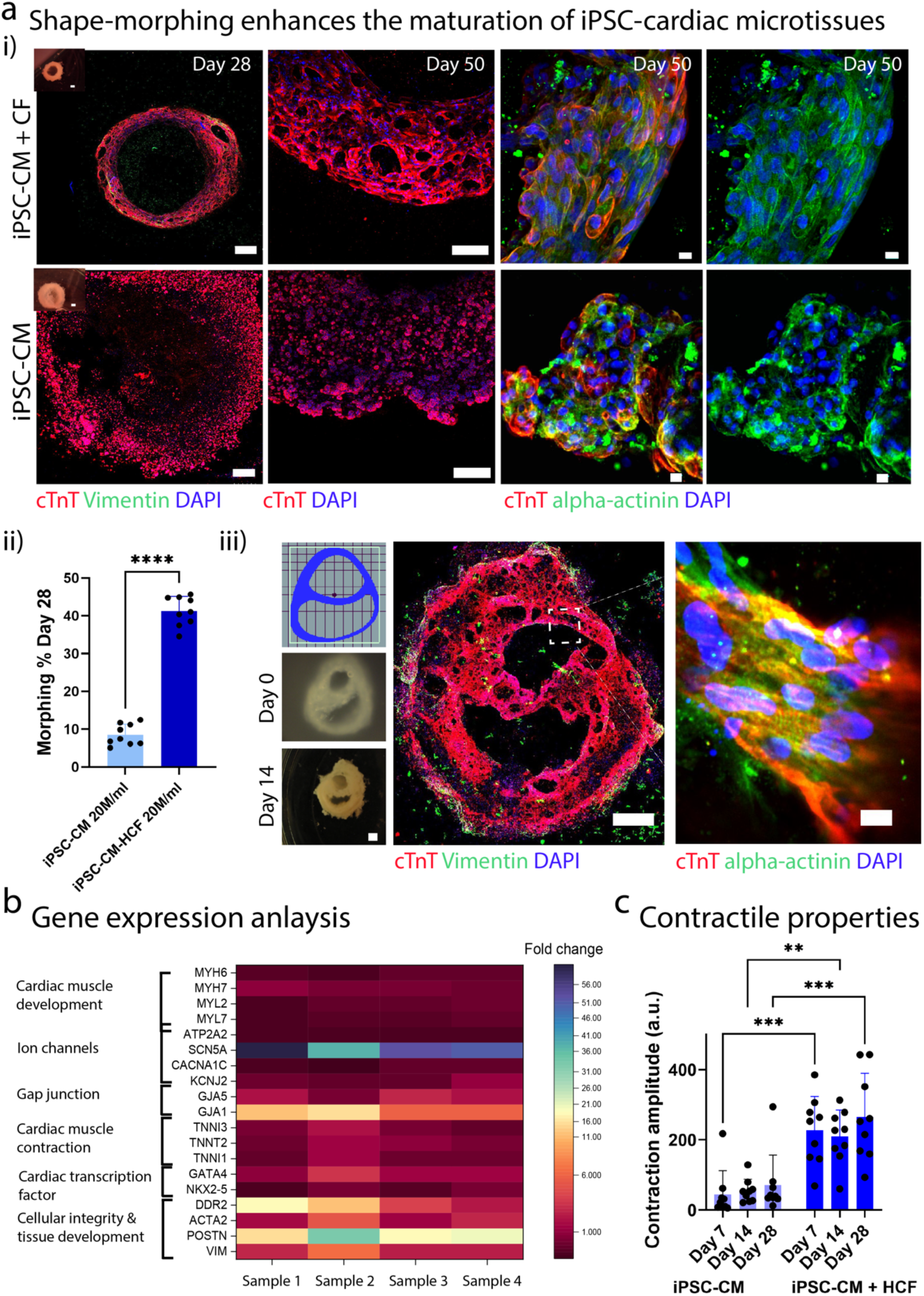
Structural and functional maturation of bioprinted iPSC-derived heart tissue: **(a)** Shape-morphing of high cell density bioprinted heart tubes containing iPSC-CMs + CFs (7:3 ratio) compared to iPSC-CMs only controls demonstrated by confocal imaging of (i) cardiac troponin T (cTnT) and vimentin at day 28 (scale bar 500µm), and (ii) cTnT and alpha-actinin at day 50 (scale bar 10µm). Inset brightfield images have a scale bar of 1mm. (iii) Quantification of shape-morphing in bioprinted heart tubes at day 28 (% diameter change), based on brightfield images (n=9 biological replicates, unpaired t-test, **** denotes p < 0.0001). Images are representative of n=3 biological replicates. (iv) Bioprinting of double ventricle heart tissue: design file (top left), and brightfield images at day 0 and day 14 (bottom left: Scale bar 1mm). Confocal image (right) demonstrating cTnT organisation at day 14 (scale bar 1000µm), and sarcomeric alpha-actinin organisation (scale bar 10µm). **(b)** Gene expression analysis of bioprinted heart tubes containing iPSC-CMs + CFs relative to iPSC-CM only controls at day 30. Fold change expression with GAPDH as the housekeeping gene (n=4 biological replicates). **(c)** Contractility analysis of shape-morphing heart tubes (iPSC-CMs + CF) compared to iPSC-CM only controls at day 7, 14, and 28 (n=9 biological replicates, two-way ANOVA with Tukey’s multiple comparison test where ** denotes p < 0.01, and *** denotes p < 0.001).

Notably, robust circumferential alignment and sarcomere maturation were evident after 50 days of culture, and importantly this structural organisation was present throughout the entire bioprinted tube (Figure 5a ii, Figure S10a ii). In contrast, iPSC-CMs in fibroblast-free controls remained rounded and disorganised, with no evidence of sarcomere maturation after 50 days (Figure 5a ii, Figure S10a ii). These findings demonstrate the CF-generated contraction forces and/or CM-CF coupling can guide the structural organisation and maturation of iPSC-CMs in shape-morphing heart tissues. To validate the scalability and adaptability of this morphing-guided maturation approach, we bioprinted larger heart tissue constructs mimicking a cross-section of the human heart using co-cultures of iPSC-CMs and CFs (Figure 5a iv, Movie S12). Mirroring the results observed in the bioprinted tubes, immunofluorescence staining revealed the emergence of cell alignment and sarcomere development along the principal tissue axes of the myocardium wall (Figure 5a iv, Figure S10a iii). Importantly, we also observed that the ratio of iPSC-CMs to CFs remained stable during culture, as evidenced by robust staining for cTnT relative to vimentin (Fig 5a i, iii). Taken together, these results support the feasibility of scaling this shape-morphing-guided maturation approach to larger and more complex tissue geometries.

To validate our findings at the transcriptional level, we performed real-time qPCR to assess the expression of genes involved in cardiac tissue development and maturation. Within this gene panel, several genes showed upregulation in shape-morphing heart tissues compared to static fibroblast-free controls at day 30 of culture (Figure 5b). Notably, we observed higher expression of genes related to electrophysiology and conduction such as *SCN5A* (encoding sodium channels, fold change 50), *GJA1* (encoding connexin 43 gap junction, fold change 9), and *GJA5* (connexin 40, fold change 1.6). The upregulation of *GJA1* and *GJA5*, both ventricular-specific markers ^43^ suggest a ventricular-like CM phenotype. Additionally, *GATA4*, a critical regulatory gene for cardiac muscle development, was also upregulated in the morphing tubes (fold change 1.6). As expected, genes related to presence of cardiac fibroblasts were upregulated in the shape-morphing heart tubes, including *POSTN* (encodes the ECM protein periostin, fold change 21.8), *VIM* (encoding vimentin, fold change 3.2), *ACTA2* (encoding alpha-smooth muscle actin, fold change 2.6), and *DDR2* (encoding discoidin domain-containing receptor 2, involved in cell-ECM interactions, fold change 8.9). Interestingly, despite observing improved structural organisation of iPSC-CMs via immunofluorescence staining (Figure 5a), we did not detect upregulation of genes directly related to cardiac muscle development or contraction at day 30. This may be attributed to the dynamic nature of gene expression during development and differentiation, as well as the limitations of assessing gene expression at a single time point. Nevertheless, the upregulation of genes associated with electrophysiology and conduction, coupled with the enhanced structural organisation of iPSC-CMs observed via immunofluorescence, strongly suggests that fibroblast-mediated 4D shape-morphing promotes the structural and functional maturation of bioprinted heart tissues.

Next, we recorded contraction videos of the bioprinted heart tubes (Movie S9, S10, S11) and employed the open-source software Musclemotion to quantify both the contraction amplitude (correlated to the force of contraction) and peak-to-peak time (a measure of beating rate) ^44^. Bioprinted tubes undergoing shape-morphing (+CFs) displayed significant four to fivefold increases in contraction amplitude compared to the iPSC-CM-only controls (Figure 5c). Notably, control samples displayed weaker, asynchronous contraction, localized to specific regions of the tissue. In contrast, shape-morphing tubes displayed synchronised contractions, starting approximately 3 days after bioprinting. Additionally, the beating rate of shape-morphing heart tissues was faster (indicated by shorter peak-to-peak times) at earlier time points, reaching approximately 950 ms by day 14 compared to 1400 ms in controls (Figure S10b).

Our results collectively demonstrate that fibroblast-mediated 4D shape-morphing can enhance the structural and functional properties of bioprinted iPSC-derived heart tissues. While previous studies have demonstrated that co-culture with CFs can improve the contractile properties of iPSC-CMs in spheroid models ^45–47^, our work reveals that fibroblast-generated contractile forces create strain fields in morphing tissues which guides the collective alignment and structural maturation of iPSC-CMs. Notably, we demonstrate that this shape-morphing-guided maturation can orchestrate collective cell organisation across substantial length scales, achieving consistent circumferential alignment across the entire 24mm circumference of our bioprinted heart tubes (Figure 5a i). Furthermore, our results show that cell-generated forces within shape-morphing heart tissues can provide endogenous mechanical stimulation that promotes tissue maturation. While exogenous mechanical stimulation via bioreactors has been shown to enhance engineered heart tissue maturation ^48^, such methods are challenging to implement for complex anatomical geometries. In contrast, our approach, leveraging bioprinting within granular support hydrogels that provide a dynamic viscoelastic environment for tissue shape evolution, is readily adaptable to diverse tissue geometries, as exemplified by our successful scaling to multi-chamber heart models (Figure 5a iv). This platform, coupled with our ability to predict the evolution of tissue-scale strain fields using FE modelling, holds great potential for bioprinting tissues that recapitulate the intricate structural anisotropies found in the human heart.

### 2.6 4D shape-morphing in granular support hydrogels: A novel framework for the design and fabrication of bioprinted tissues with advanced maturation

We introduce a novel 4D bioprinting approach where tissues are programmed to undergo shape-morphing driven by endogenous cell-generated forces, ultimately sculpting their final form, composition, and function. Our platform utilises soft granular support hydrogels with shear-thinning and self-healing properties, enabling them to: 1) receive tissues with defined geometries via embedded printing, and 2) support the structural evolution of the tissue post-printing. Notably, we discovered that the support hydrogel viscoelasticity can actively influence cell and ECM assembly within the shape-morphing tissues by providing a mechanical constraint during their structural evolution. While support hydrogels for bioprinting have been traditionally viewed as structural supports to prevent bioink collapse, our results highlight how their properties can be modulated to guide post-printing tissue maturation, thereby expanding their utility in biofabrication. A key advantage of our approach is the ability to design and predict 4D shape-morphing behaviours through a combination of bioprinting and FE modelling. The initial geometry of the tissue, precisely controlled through embedded bioprinting, guides its structural evolution, which is accurately predicted by our FE model (Figure 6a). This platform opens up exciting possibilities for bioprinting tissues that undergo more complex shape-morphing behaviours observed during organ morphogenesis. Furthermore, we observed that 4D shape-morphing significantly enhanced cell and ECM organisation and maturation within our bioprinted tissues, which is a major challenge in the field ^13^. Our advanced FE models helped elucidate the underlying principles of this behaviour, revealing how active cell stresses, stress fiber remodelling, and collagen fiber remodelling generate principal strain directions that guide cell alignment via the principle of stress avoidance (Figure 6b). This generalisable understanding can help enable application of these principles for a wide range of 4D bioprinting applications.

**Figure 6.**
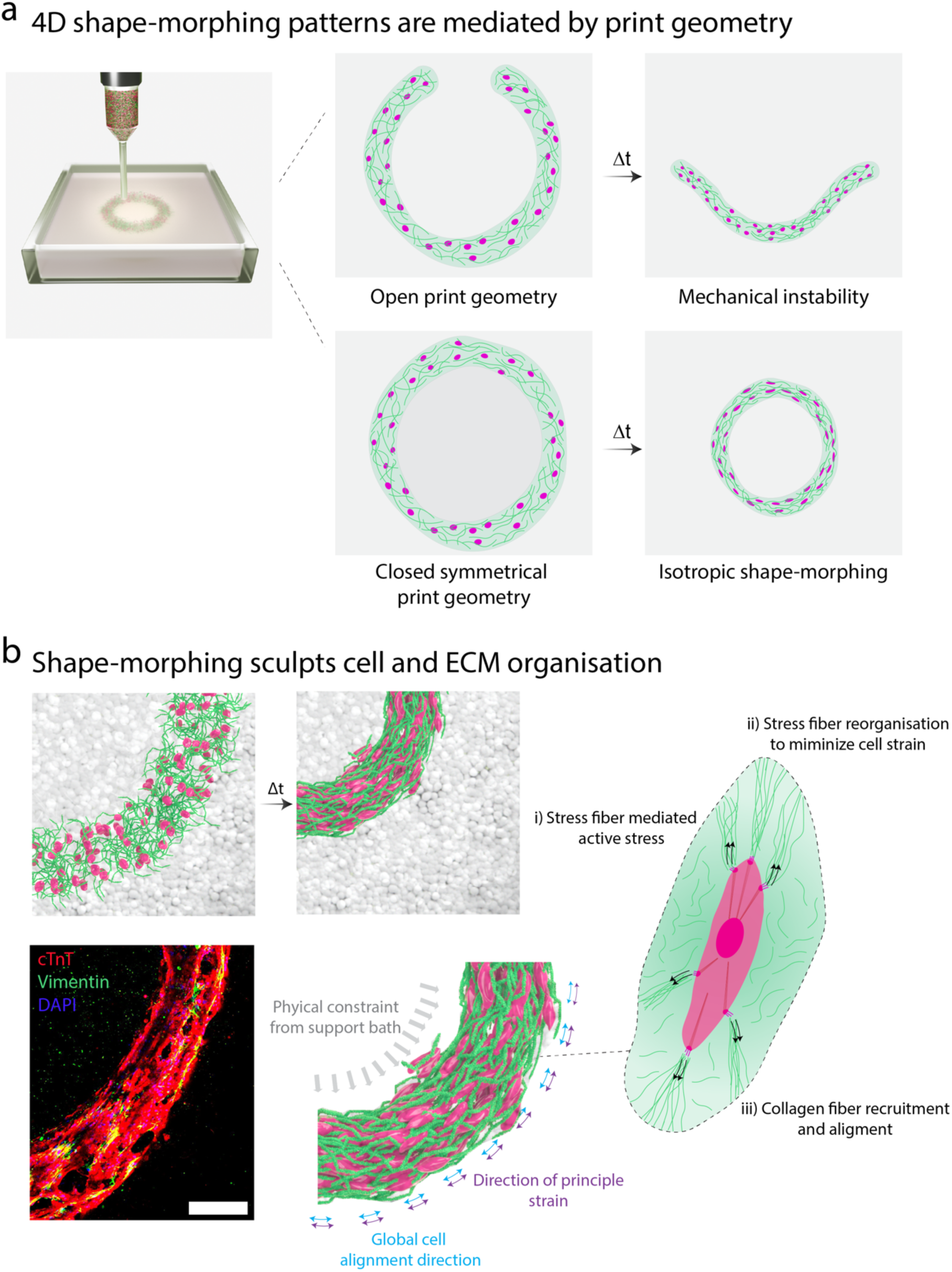
Overview of 4D bioprinting platform where tissues are programmed to undergo shape-morphing to sculpt their maturation: **(a)** The initial print geometry influences the shape-morphing patterns that emerge under cell-generated contraction forces. Open geometries exhibit mechanical instabilities, while closed symmetrical geometries undergo isotropic shrinkage. **(b)** As cells exert active contractile forces on the collagen network printed within the support bath, they experience physical resistance which drives stress fiber remodelling and collagen fiber remodelling to minimise cell strain. This leads to cell alignment along the principal tissue axis (i.e. the direction of maximum stretch). Additionally, the viscoelasticity of the support hydrogel provides physical resistance to shape-morphing, which enhances the extent of cell reorganisation. This 4D shape-morphing-guided remodelling can be harnessed to enhance the maturation of bioprinted iPSC-derived heart tissues (scale bar 100µm).

## 3. Conclusion

In conclusion, our novel bioprinting platform facilitates the fabrication of tissues capable of programmable and predictable 4D shape-morphing driven by cell-generated forces. We demonstrate that 4D tissue shape-morphing can sculpt changes in internal cell and ECM alignment, offering a platform for programming anisotropy and maturation in bioprinted constructs. Crucially, it was possible to model these behaviours at the cell and tissue scales using thermodynamically-motivated models for stress fiber and collagen fiber remodelling. We anticipate that this ability to design, predict, and program 4D shape-morphing in bioprinted tissues will open up numerous opportunities for engineering organ rudiments that recapitulate morphogenetic processes to sculpt their final form, composition, and function. This offers new directions in organ bioprinting focused on recapitulating developmental processes rather than the end-stage geometrical structure of the organ.

## 4. Experimental Section

### Primary Human cardiac fibroblasts (HCF) culture

Primary human cardiac fibroblasts (C-12375, Promocell) were thawed and maintained in Fibroblast growth medium 3 (C-23025, Promocell) for 2-3 passages, followed by expansion in Gibco™ MEM alpha-GlutaMAX^TM^ medium (32561037, Fisher Scientific) supplemented with 10% fetal bovine serum (FBS), 1% penicillin-streptomycin and 5ng/ml Human Recombinant Fibroblast Growth factor-2 (FGF-2) (130093564, Miltenyi Biotec). Fibroblasts were passaged using trypsin EDTA 0.5%, and the centrifuged at 300G for 5 minutes, followed by mixing the cells with hydrogel-based ink for 3D culture and bioprinting. The cells were expanded between 7-15 passages for 3D culture and bioprinting experiments. Further details of all cells used can be found in Table S1.

### Primary Human umbilical vein endothelial cell (HUVEC) culture

Primary HUVECs (C-12200, Promocell) were thawed and maintained in endothelial growth medium (C-22010) supplemented with 1% penicillin-streptomycin. HUVECs were expanded until they reached 90% confluency and then, dissociated using trypsin EDTA 0.5% for 5mins incubation, followed by centrifugation at 300G for 5mins. 3D culture and bioprinting experiments were performed using cells between passages 4-7.

### Human induced pluripotent stem cell culture (iPSC)-maintenance and expansion

Human Induced Pluripotent Stem cells (A18945, Fisher Scientific) were seeded on Corning^TM^ Matrigel (354277, Fisher Scientific) coated six-well plates. The Matrigel coating solution was prepared by diluting Matrigel (2%) in Gibco^TM^ DMEM F-12 GlutaMAX™ supplement medium (10565018, Fisher Scientific), and the plates were coated for 1 hour at room temperature, followed by incubation at 37°C for 20mins. iPSCs were thawed using 5µM Rock Inhibitor Y-27632 (72304, STEMCELL Technologies) that was added in the culture medium to enhance cell survival. To detach the cells, iPSCs were passaged using Accutase^TM^ (07920, or 07922, STEMCELL Technologies) for 5 minutes in incubation at 37°C. Detached cells were centrifuged at 200G for 3 minutes, and the cell pellet was then gently dissociated to seed them either on coated well plates or embedded in ink. iPSCs were maintained in mTeSR^TM^ supplemented (85857, STEMCELL Technologies) medium, mTeSR^TM^ Plus (100-1130, STEMCELL Technologies) or Gibco^TM^ Essential 8^TM^ medium (A2858501, Fisher Scientific), which was changed daily until the cells reached 80-90% confluency (typically 3-4 days). To perform iPSC bioprinting, the cells were used between passages 5-8. Rock inhibitor (10*μ*m) was added to the medium for 24h post-bioprinting to enhance cell survival and then replaced with fresh medium.

### iPSC differentiation to cardiomyocytes and purification

iPSCs were cultured on matrigel-coated 12 well plates, and differentiation was initiated when the cells reached 65-80% confluency using the well-established GIWI protocol ^49,50^. For the first 24 hours, cells were maintained in Gibco™ RPMI 1640 medium (11875085, Fisher scientific) supplemented with Gibco™ B27 minus insulin (2%) (A1895601, Fisher Scientific) and 10µM CHIR99021 (72054, Stemcell technologies). Following this, the media was changed to RPMI 1640 medium supplemented with 3-5µM CHIR99021 (Wnt signalling pathway activation) for 48h. Then, the media was switched to RPMI 1640 medium supplemented with 5µM IWP-2 (72124, Stemcell technologies) to inhibit the Wnt pathway with daily media changes for another 48h. On day 8, the culture medium was changed to RPMI 1640 supplemented with Gibco™ B27 supplement (2%) (17504044, Fisher Scientific) and changed after every 2 days until day 14.

We then employed a well-established metabolism-based purification protocol to enrich the cardiomyocyte population^51^. The purification protocol was initiated on day 14 by changing the culture media to Gibco™ RPMI 1640 Medium, no glucose (11879020, Fisher Scientific) supplemented with 1% penicillin-streptomycin, 4mM Sodium Lactate (L7022, Merck) and 0.5mM L-ascorbic acid 2-phosphate (A8960, Merck). Media changes were performed every two days. Following 4 days in purification media, the culture media was changed to Gibco™ MEM alpha-GlutaMAX^TM^ medium (32561037, Fisher Scientific) supplemented with 10% fetal bovine serum (FBS), 1% penicillin-streptomycin and 200µM L-ascorbic acid 2-phosphate (maintenance medium: CM medium) for cardiomyocyte maintenance^52^. For subsequent bioprinting experiments, the iPSC-cardiomyocytes were detached through incubation in Gibco™ TrypLE™ Select Enzyme 10X (A1217701, Fisher Scientific) for 20mins, followed by centrifugation at 200G for 3mins. The iPSC-cardiomyocytes were gently dissociated into single cells and then added to a collagen bioink. For first 24h post-bioprinting, rock inhibitor (10µM) was added to the maintenance medium to promote cell survival.

### Bioink preparation

Type I bovine atelocollagen (PureCol®3mg/ml, Nutragen®6mg/ml, FibriCol®10mg/ml from Advanced Biomatrix) were neutralized as per the manufacturer’s instructions. Briefly, one part of chilled 10X PBS was first added to eight parts of chilled collagen solution and then 0.2M NaOH were added to adjust the pH to around 7.2. The remaining volume was adjusted to 10 parts with cell suspension medium to a final concentration of 2.4mg/ml, 4.8mg/ml and 8mg/ml, respectively. Sodium hyaluronate (HA) 4wt% (50-90kDa) (6000219, Contipro-Czech Republic) was added to the collagen solution to enhance the viscosity for printability. The bioink was then kept in ice box to prevent gelation before bioprinting. For experiments involving iPSCs, 5% volume matrigel (dilution factor 310-320ul) was added to the original bioink (collagen I 2.4mg/ml, HA 40mg/ml & Matrigel 5%) and for the cell type studies (Figure S3) to maintain consistency between groups. We mixed a fluorescently tagged collagen (C4361, Merck) with untagged collagen 3mg/ml to visualise the cellular interactions with collagen fibres and their alignment. The final collagen concentration was 1.9mg/ml (232ul of FITC-labelled collagen with 568ul PureCol®3mg/ml) after neutralization with 0.2M NaOH, and cell medium was added to adjust the volume to 1ml. The complete bioink with HA was later crosslinked at 37°C incubation for 45 minutes post-bioprinting to crosslink the collagen network.

### Agarose support bath fabrication

Agarose type I, low EEO was purchased from Sigma-Aldrich (A6013). Agarose microparticles were prepared using a shearing technique to form the viscoelastic suspension bath with varied stiffness ^53^. Briefly, a 0.5wt% agarose solution (1X PBS) was autoclaved at 121°C, and then the molten solution was allowed to cool at room temperature for 3 hours under stirring at 700rpm. This protocol results in the formation of microparticles via shearing as the agarose solidifies. The agarose microparticle solution was maintained in sterile conditions and stored for up to 3 months at 4-7°C. To reduce the stiffness and packing density of microparticles, agarose solution was diluted in PBS to prepare 60% and 80% particle concentrations.

### Rheological characterization

The rheological properties of the collagen ink and collagen-HA ink were characterised using rotational shear rheometry (25mm diameter, gap 1mm; Anton Paar MCR 302) at 25 °C. To assess the shear-thinning properties of bioink, viscosity was measured as a function of shear rate (0–100 s^−1^). Storage and loss modulus were measured as a function of shear strain (0.0001-1). Thixotropic properties of the agarose support bath were assessed by applying low (1 s^-1^) and high (100 s^-1^) stress rate, periodically.

### 3D bioprinting setup and embedded bioprinting

Designs for bioprinting were prepared using Autodesk Inventor and then converted into g-code using Repetier-host software where key process parameters such as the printing speed (2-7mm/s) and infill patterns (concentric & rectilinear) were adjusted. Bioprinting experiments were performed using an Allevi 2 bioprinter placed inside a biosafety hood to maintain sterile conditions. The pneumatic extrusion pressure varied depending on the printing nozzle, collagen concentration and geometry printed. We employed 4-10 psi for 30G needles in HCF and cell type experiments, and 2 psi for 25G needles during high-density iPSC-CM bioprinting. All bioprinting was performed at 22°C, and post-bioprinting, constructs were incubated immediately at 37°C for physical crosslinking.

High glass bottom µ-Dishes (35 mm, 81158, ibidi) were used for culturing the bioprinted constructs. These dishes enabled imaging of the tissue within the support bath during the culture period using inverted microscopy. The culture dishes were adapted to contain two compartments separated by a solid agarose divider (2wt%). Culture medium was added in one compartment and the agarose suspension bath was added to the second compartment ^54^. The agarose divider allowed the diffusion of culture medium into the bioprinted constructs suspended in the support bath. The bioink was extruded into agarose microparticle as support bath, followed by incubation at 37°C for physical cross-linking. The agarose suspension bath supported the bioprinted tissue for the first 7 days of culture and was then gradually removed and diluted by media exchanges.

Bioprinted constructs containing fibroblasts were cultured in MEM alpha-GlutaMAX^TM^ medium supplemented with 10% FBS, 1% penicillin-streptomycin and 5ng/ml Human Recombinant Fibroblast Growth factor-2. Bioprinted constructs containing iPSCs and HUVECs were cultured in mTESR and endothelial growth medium respectively with 1% penicillin-streptomycin. Co-printed constructs with iPSCs/HCF, or HUVECs/HCF (Figure. 3) were cultured in respective standard medium, not HCF medium.

### Design of experiments (DOE)

To study the effect of cell density and collagen concentration on tissue shape-morphing, the DOE study was planned based on a full-factorial method using Minitab software (Figure 3a i). The HCF cell density and collagen concentration varied from 0.5 to 10 million/ml and 2.4-8mg/ml respectively, keeping the HA concentration (4wt%) constant. All the bioprinting experiments were based on the tube geometry of 20 layers, 7mm/s printing speed and 4-10psi (depending on collagen concentration) at 22°C. The morphing percentage (reduction in diameter of the tube) was plotted against the cell density and collagen concentration in a three-dimensional surface plot using Minitab.

### Microscopy analysis of bioprinted constructs

To visualize live samples in the culture period, a brightfield microscope (Dinocapture 2.0) was used. Images of the bioprinted constructs were captured at different time points and FIJI software was used to measure shape change relative to day 0. Brightfield images were converted to coloured brightfield using ‘Royal’ lookup option in FIJI software. Inverted confocal microscopy was also used to image bioprinted samples containing FITC-conjugated collagen and cell dye. Invitrogen^TM^ CellTracker™ dyes (C34552/C7025, Fisher Scientific) were used to track the cells. Bioprinted tissues were visualized live on the day of bioprinting (day 0) and the terminal day of culture using multi-channel confocal microscopy (Olympus FV-1000/3000).

### Live/dead staining

Live/dead staining was performed by treating samples with 2µM Calcein-AM (11564257 Fisher Scientific) and 4µM Ethidium homodimer (E1903, Merck) in 1X PBS, followed by 1-hour incubation at 37°C. The samples were imaged using an FV1000 confocal microscope.

### Actin-myosin inhibition study

To understand the shape-morphing mechanisms within the bioprinted constructs, cells were treated with potential inhibitors such as rock inhibitor Y-27632 (10µM), blebbistatin (20µM), and cytochalasin D (10µM). The inhibitors were added to MEM alpha-GlutaMAX^TM^ medium supplemented with 10% FBS and 1% Penicillin streptomycin, and cells were treated with the respective inhibitor for 30 minutes before mixing with bioink for bioprinting. The bioprinted constructs were cultured in the inhibitor-treated medium for seven days. The extent of shape-morphing (reduction in diameter of tubes relative to day 0) was then assessed using brightfield imaging (Dinocapture 2.0) and FIJI software.

### Immunofluorescent staining

Samples for immunofluorescent staining were fixed with 10% neutral buffered formalin solution overnight at 4°C and stored in DPBS the following day until staining. Samples were then permeabilized with Triton-X 100 (0.2%) in PBS and 2% BSA for 1h and then blocked using 2% BSA in PBS for 2h. Primary antibody (1:200 dilution in blocking buffer) staining was then carried out overnight at 4°C, followed by three times washing with PBS and then, adding the secondary antibody for 2 hours (1:200 dilution in blocking buffer).

Samples were washed thrice before being counterstained with Fluoroshield^TM^ DAPI (F6057, Merck) for 20mins. Imaging was performed in high-resolution confocal microscopy. All primary and secondary antibodies are listed in the reagent list below (Table S2).

### Phalloidin TRITC staining

To visualize the actin cytoskeleton, fixed bioprinted constructs were stained with 0.05mg/ml phalloidin-tetramethylrhodamine B isothiocyanate (P1951, Merck) overnight at 4°C. The constructs were then washed twice with 1X PBS followed by staining with Fluoroshield^TM^ DAPI for 20mins at room temperature. The samples were visualized in confocal microscopy.

### Nanoindentation

Mechanical properties of bioprinted samples were quantified using a Optics11 Life Pavone nanoindenter using 29.5 µm tip radius and 4.18 N/m stiffness probe in air. We compared the indentation modulus of the day 0 sample with day 14 bioprinted sheet tissue (2M/ml HCF in 2mg/ml collagen and 4wt% HA).

### Quantifying cell and collagen fibre alignment

Cell and collagen fibre alignment analysis was performed by measuring the probability distributions of cells and collagen fibres at a particular angle relative to the printing direction. Confocal imaging of the bioprinted tissue was used to image the cells (stained with rhodamine-phalloidin) and collagen fibres (FITC-conjugated). The multicolored images were split into separate channels, i.e. red for cells and green for FITC-collagen. The coloured images were then converted to greyscale images, and the orientation data was extracted using the directionality plugin in FIJI (Image-J) selecting angle 0-180° ^36,55^. This orientation data was then plotted in polar histogram format using ORIGIN 2023 software. The greyscale images were also converted into colour survey maps using the OrientationJ plugin in FIJI, where each colour corresponds to a particular direction relative to the principal axis demonstrating the alignment.

### Contraction amplitude and peak-to-peak time analysis

Brightfield videos of bioprinted iPSC-cardiac tissue were captured using inverted microscopy (Leica DMi; 10x magnification) for 30s at different timepoints. Videos were recorded at room temperature. The contraction amplitude and peak-to-peak time were analysed using the FIJI Musclemotion macro ^56^. Contraction graphs were plotted using GraphPad software.

### RNA isolation and PCR array

TRIzol® reagent (15596018, Invitrogen™) and choloform (C2432, Merck) were used with PureLink™ RNA Micro Kit (12183016, Invitrogen™) to isolate the RNA from bioprinted iPSC-CM samples with and without HCF on day 30 (20M/ml cell density). TRIzol® reagent extracted the RNA with choloform for phase separation, and the extracted RNA was washed, purified following the micro kit instructions. cDNA preparation was carried out using the RT^2^ First Strand Kit (330404, Qiagen) and processed in veriti Thermal cycler. This was followed by the addition of the cDNA and RT^2^ SYBR Green ROX qPCR Mastermix (330522, Qiagen) to a custom designed RT2 PCR Array 96-well plate (330171, Qiagen) maintaining 10% excess amount for qPCR run. QuantStudio 5 was used to run the qPCR, and data analysis was performed in excel using the C^T^ value. Heatmaps were plotted in OriginPro 2023 software considering GAPDH as the housekeeping gene, and bioprinted iPSC-CM tubes as the control sample. Further details on the genes assessed can be found in Table S3.

### Statistical analysis

All experimental data was compiled and stored in Microsoft Excel. All graphs are presented as mean with standard deviation along with sample numbers denoted in each graph and figure legend. The reported sample sizes (n-numbers) in the figure legends denote biologically independent samples throughout. Statistical analysis was performed using GraphPad Prism 10 software. Two-way or one-way ANOVA tests were used depending on the number of independent variables within the experiment and Tukey’s multiple comparison tests were used to compare differences between means. P values are described as follows, ns denotes not significant, and * denotes p< 0.05, ** denotes p<0.01, *** denotes p< 0.001, and **** denotes p< 0.0001.

### Computational modelling

Briefly, we consider the formation of stress fibres of concentration *η̂_i_* within cells in a number of discrete directions *n_θ_*, with full details provided in Supplementary Section S2. The active Cauchy stress tensor ***σ***^*act*^ is then given by

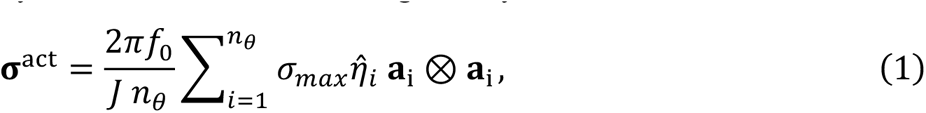

where *σ_max_* is the maximum isometric stress in a SF, *f*_0_ is the combined volume fraction of fibroblast cells and their cytoskeletal proteins, *J* is the determinant of the deformation gradient **F**, and **a**_i_ = **Fa**_i,0_ where **a**_i,0_ is a unit vector in direction *i*. The passive hyperelastic behaviour of the collagen matrix is similarly described such that

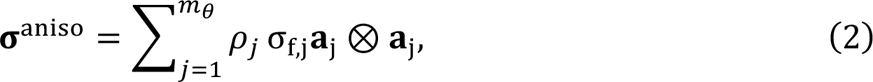

where *ρ_j_* is the fibre density in direction *i* and σ_5,i_ is the fibre bundle stress in that direction (see Supplementary Section S2). A neo-Hookean model captures the underlying isotropic behaviour of the bioink ***σ***^*iso*^, combined to describe the total bioink stress:

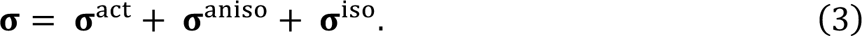

The mechanical behaviour of the support bath granular hydrogel is also described as a neo-Hookean material. This framework was implemented in finite element analysis software Abaqus (2021) through a user-defined material subroutine (UMAT). Finite element meshes of the bioprinted tissues in their reference (day 0) states were generated embedded in a support bath. The boundary dimensions of the support bath hydrogel were sufficient to avoid edge effects.

## Acknowledgements

This publication has emanated from research conducted with the financial support of the Irish Research Council GOIPG/2022/485) and the European Research Council (Grant number 101077900). This publication has also emanated from research supported in part by a grant from Science Foundation Ireland and is co-funded under the European Regional Development Fund under Grant number 13/RC/2073_P2. We would like to acknowledge the University of Galway College of Science and Engineering Postgraduate Research Scholarship Scheme. We would like to acknowledge the Centre for Microscopy and Imaging facilities at the University of Galway, and Dr Peter Owens for his support. The authors gratefully acknowledge the Genomics and Screening Core Facility, and Dr Enda O’Connell for their support & assistance with PCR analysis. We are grateful to Dr Arun Thirumaran for guidance on RNA isolation, and Dr Olena Kudina for helping with Nanoindentation experiments. We acknowledge Dr Daniel Kelly and Dr Vasileios Sergis for support with heart CAD file design, g-code optimisation, and recording of the 3D bioprinting movies. We are grateful to Prof Patrick McGarry for contributions to development of the computational model in prior publications. Finally, we thank Rahul Bhatti, Sogol Kianersi, and Eoin Ó Donovan for their assistance in optimising the 2D iPSC-differentiation protocol.

## Data availability

The data that support the findings of this study are available from the corresponding author upon reasonable request.

## Table of contents image and text

A novel 4D bioprinting platform enables fabrication of tissues that undergo predictive and programmable shape-morphing driven by cell-generated forces. This method utilises embedded bioprinting to deposit ECM bioinks into yield-stress granular hydrogels that support and regulate morphing. These 4D shape-morphing patterns are tunable by adjusting print geometry, cell type, and bioink composition, and can be predicted using a computational model.

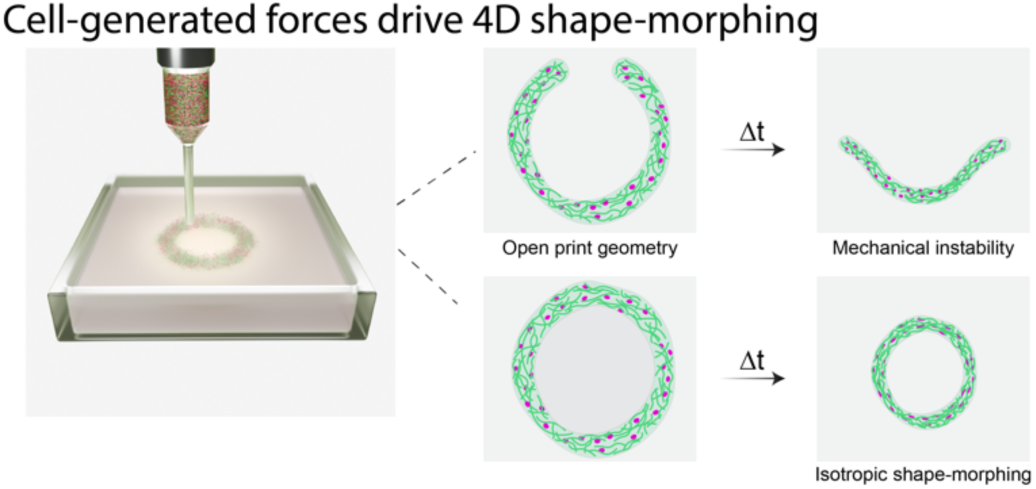

## Supporting Information S1 – Additional Figures

**Figure S1.**
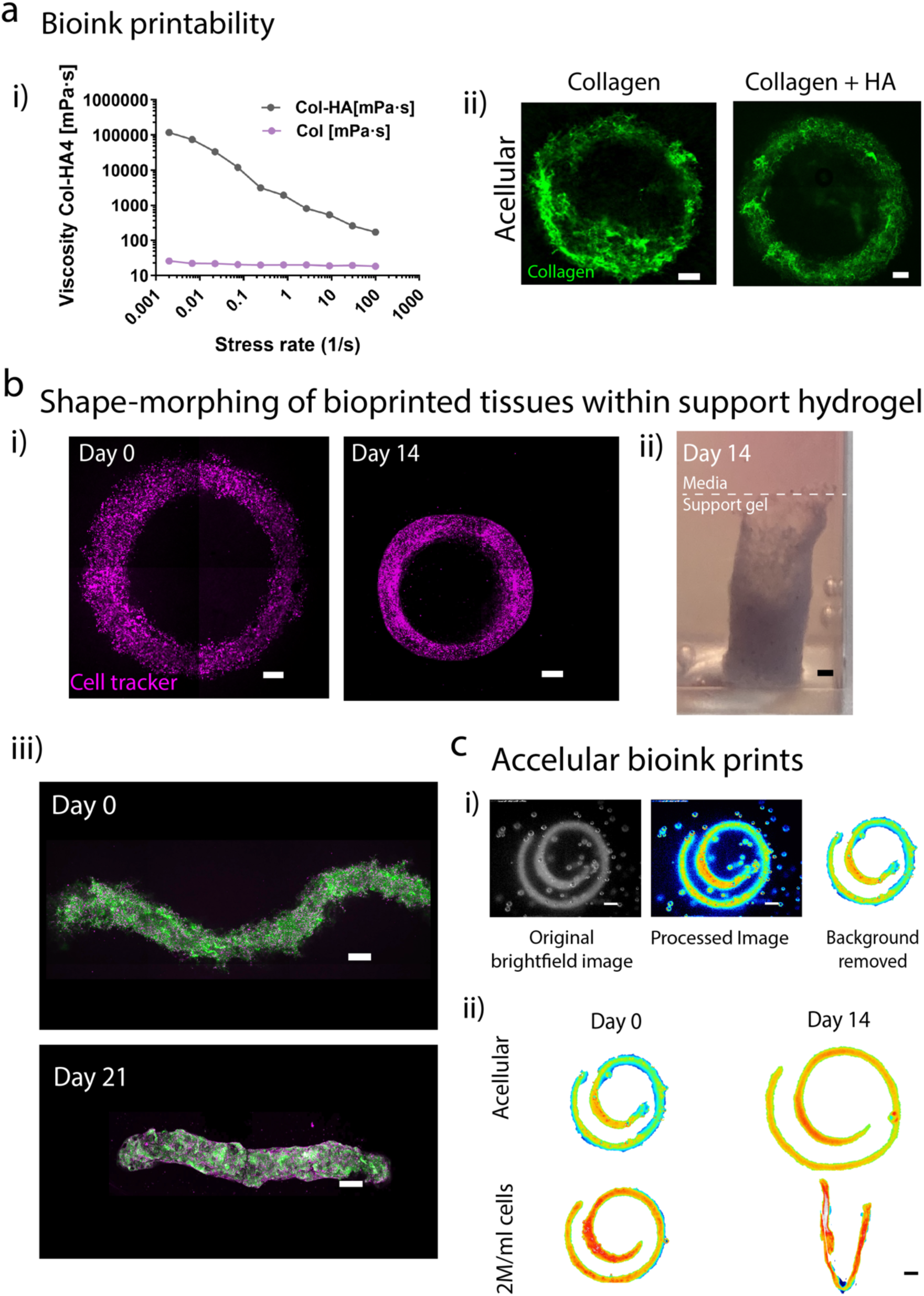
Bioink optimisation and post-bioprinting shape-morphing in granular support bath: **(a)** Hyaluronic acid (HA) improves bioink printability demonstrating through, (i) rheological characterisation of shear-thinning properties (viscosity versus stress rate) with and without HA, and (ii) confocal imaging of tubes bioprinted within the support bath, scale bar 500µm. **(b)** 4D shape-morphing of tubes bioprinted in granular support hydrogels over 14 days culture, (i) Confocal imaging of tube cross-sectional at day 0 and day14 (scale bars 500µm), and (ii) sideview of tube at day 14 (scale bar 1mm). (iii) Shape-morphing of wave geometry which unfolded over 21 days of culture within the support hydrogel (scale bar 500µm), n=3 biological replicates. **(c)** (i) Overview of brightfield image processing steps where images are processed to distinguish the bioink from the support hydrogel. (ii) Demonstration of the absence of shape-morphing in acellular bioprinted constructs (scale bar 1mm), n=3 biological replicates.

**Figure S2.**
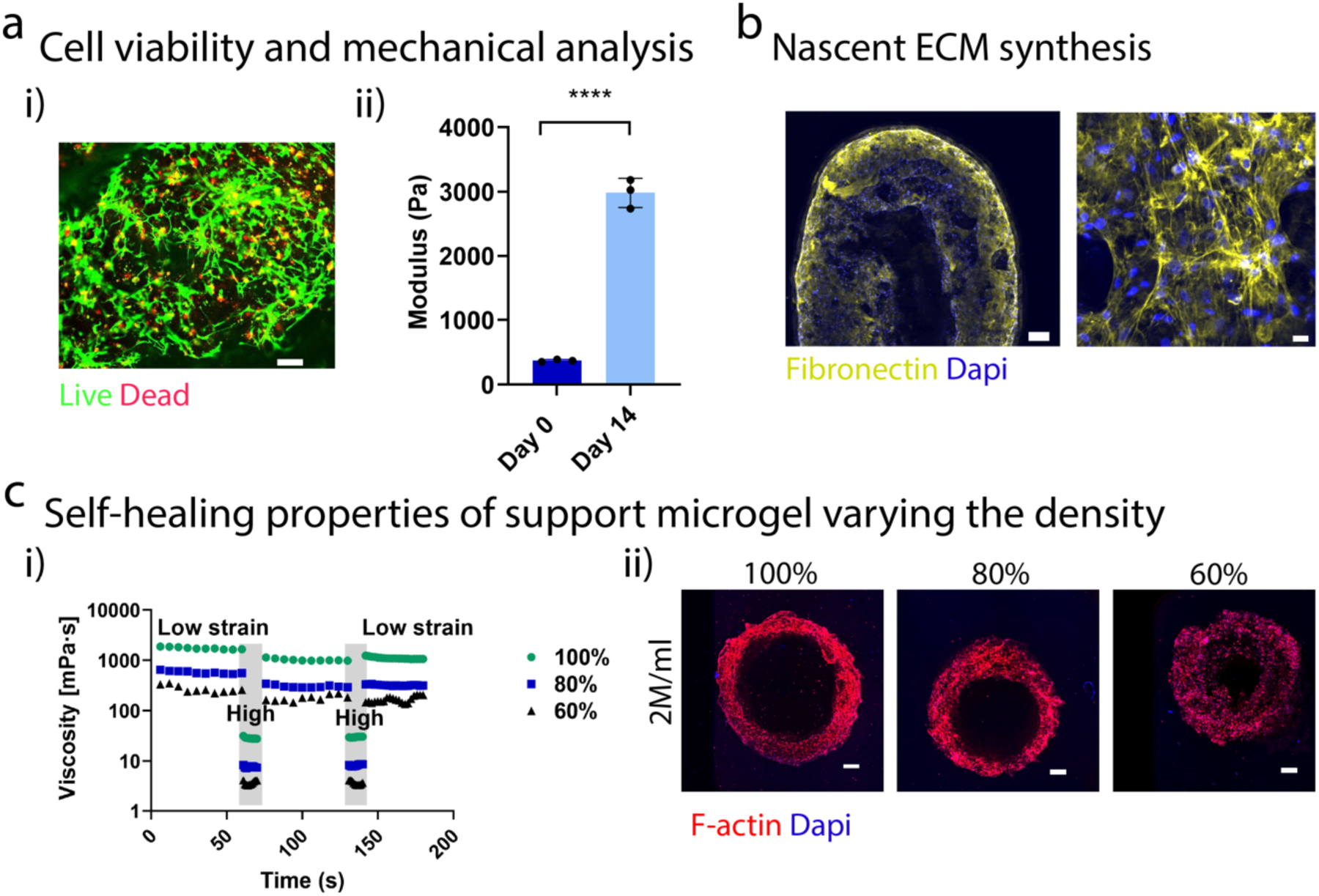
Cell viability, tissue stiffening, and nascent ECM synthesis in bioprinted tubes and influence of support bath viscoelasticity on 4D tissue shape-morphing: **(a)** (i) Live-dead staining on day 14 in shape-morphing tissues, (ii) Mechanical characterization of bioprinted morphing tissue on day 14 compared to day 0 using nanoindentation technique (n=3 biological replicates, unpaired t-test where **** denotes p <0.0001) (ii) Confirmation of nascent fibronectin secretion in bioprinted morphing tube on day 14, scale bar 200µm, 20µm (right). **(b)** (i) Influence of microgel packing density (60%, 80% and 100%) on shear thinning and self-healing properties of support hydrogel analysed through thixotropic tests, and (ii) Influence of support hydrogel viscoelasticity on tissue shape-morphing analysed through immunofluorescence staining of bioprinted tubes. Images are representative of n=3 biological replicates.

**Figure S3.**
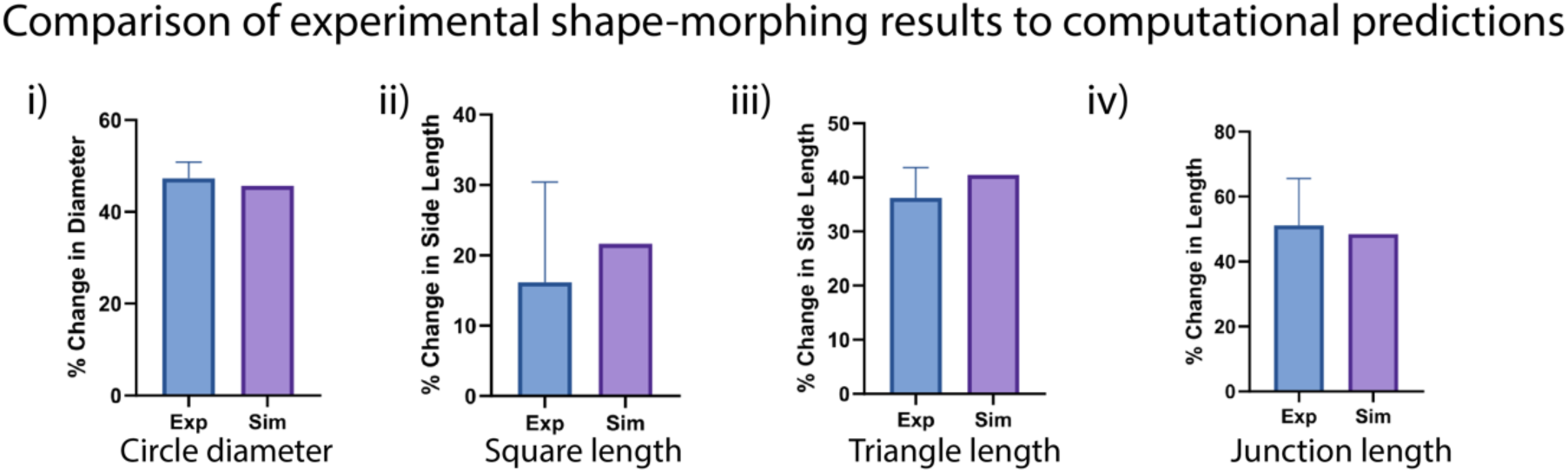
Percentage change of relevant length for each tissue geometry (circle, square, triangle, and junction) during shape-morphing from the experimental results (blue) and simulation predictions (indigo). The experimental images are presented in main figure 2. The % changes are calculated by measuring the relevant length at day 0 and day 14.

**Figure S4.**
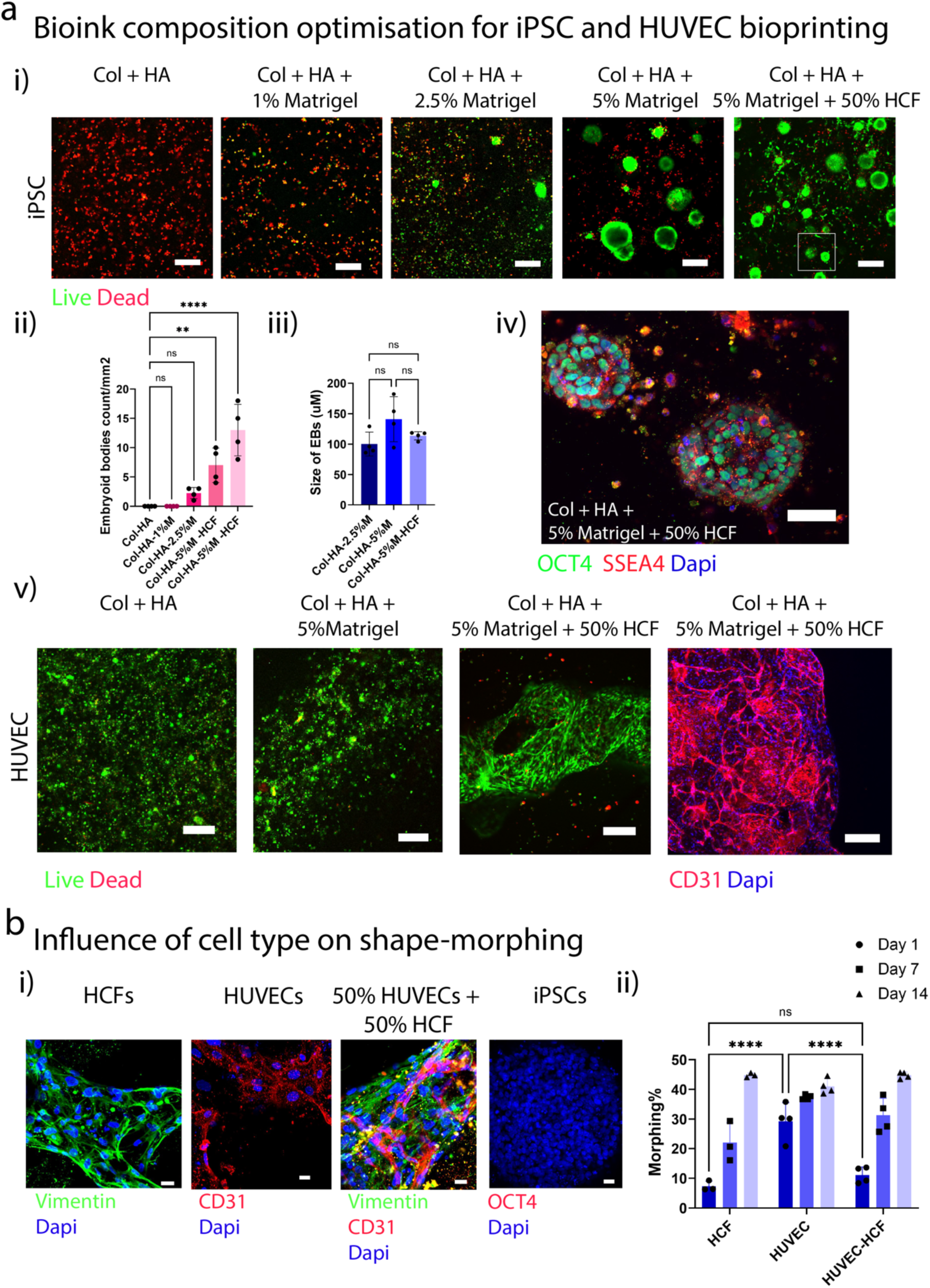
Optimising bioink composition to study the influence of cell type of 4D shape-morphing: **(a)** (i) Influence of matrigel concentration of iPSC viability in collagen hyaluronic acid bioinks analysed through live dead staining on day 7 (Scale bar 200µm). The influence of co-culture with CFs on cell viability was also assessed. (iii-iv) Evaluation of embryoid body (EB) formation in iPSC containing bioinks via quantification of (ii) EB number (n=4, biological replicates), (iii) EB size (n=10 biological replicates) and performed one-way ANOVA with Tukey’s multiple comparison test where ns denotes not significant, * denotes p < 0.05, ** denotes p < 0.01 and **** denotes p < 0.0001, and (iv) immunofluorescence staining for OCT4 (scale bar 50µm). (v) HUVEC viability in different bioink compositions analysed through live dead and CD31 staining on day 7 (scale bar 200µm). These cell encapsulation experiments presented in panel Figure S4a were performed in non-bioprinted conditions (i.e. the bioinks were cultured in standard well plates rather than the support hydrogel). **(b)** Analysis of cell phenotype in bioprinted morphing tubes after 7 days of culture characterized through (i) immunostaining for vimentin (fibroblasts), CD31 (endothelial cells), OCT4 (iPSC-pluripotency) (scale bar 20µm). (ii) Quantitative assessment of shape-morphing in tissue tubes containing HCFs, HUVECs and HCF + HUVEC (1:1 ratio) at different time points (n=3 biological replicates, Two-way ANOVA with Tukey’s multiple comparison test where ns denotes not significant, * denotes p < 0.05, ** denotes p < 0.01).

**Figure S5.**
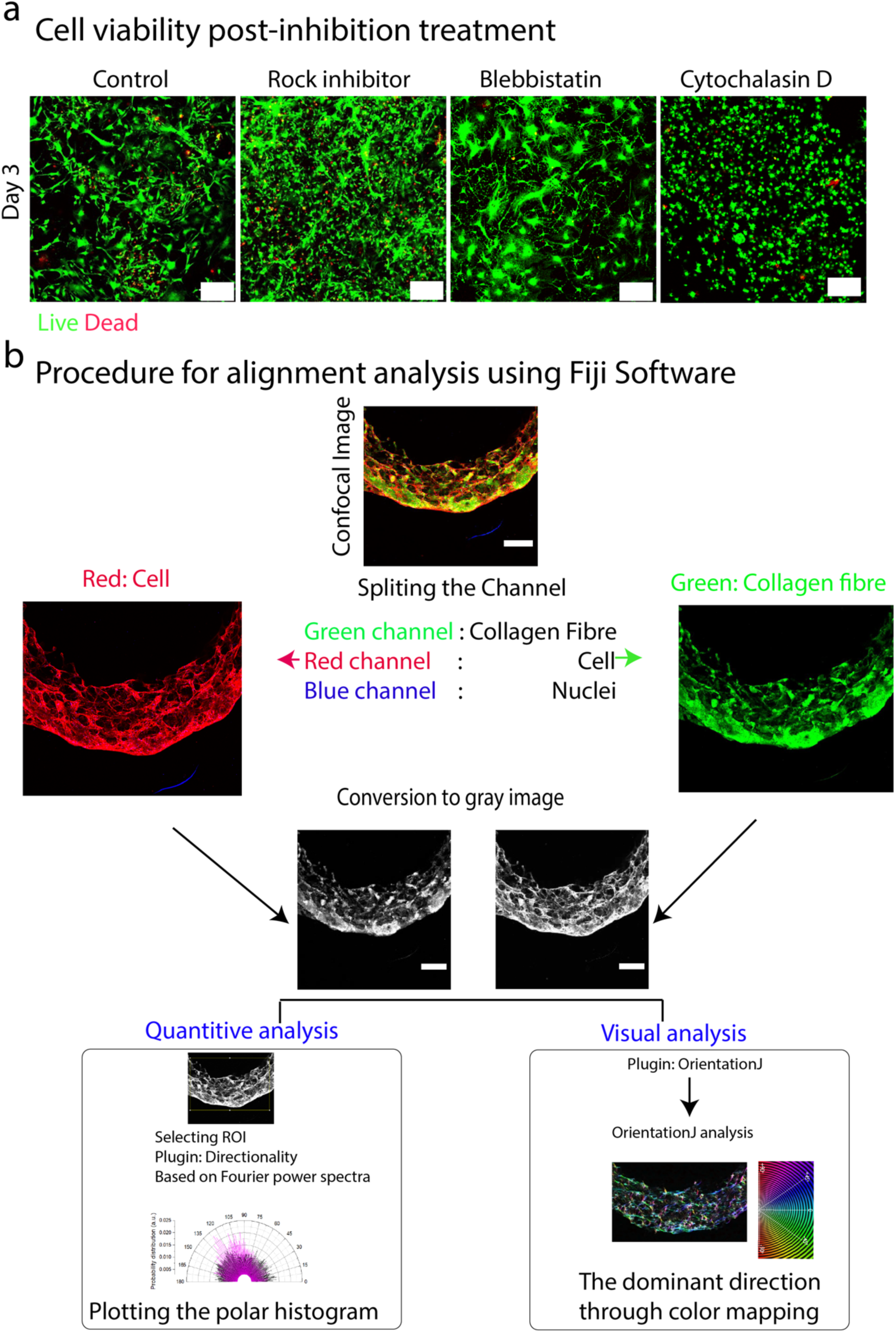
Cell viability and cell alignment analysis: **(a)** Live-dead assessment in control and inhibitor-treated morphing tissues on day 3. (Scale bar 200µm **(b)** Procedure for alignment analysis from confocal images using Fiji software. Firstly, the fluorescent channels (cells or collagen fibres) are separated followed by conversion into grayscale images. Next, the ‘directionality’ and ‘orientationJ’ plugins are used to quantify alignment and visualise the preferential direction through colour mapping.

**Figure S6.**
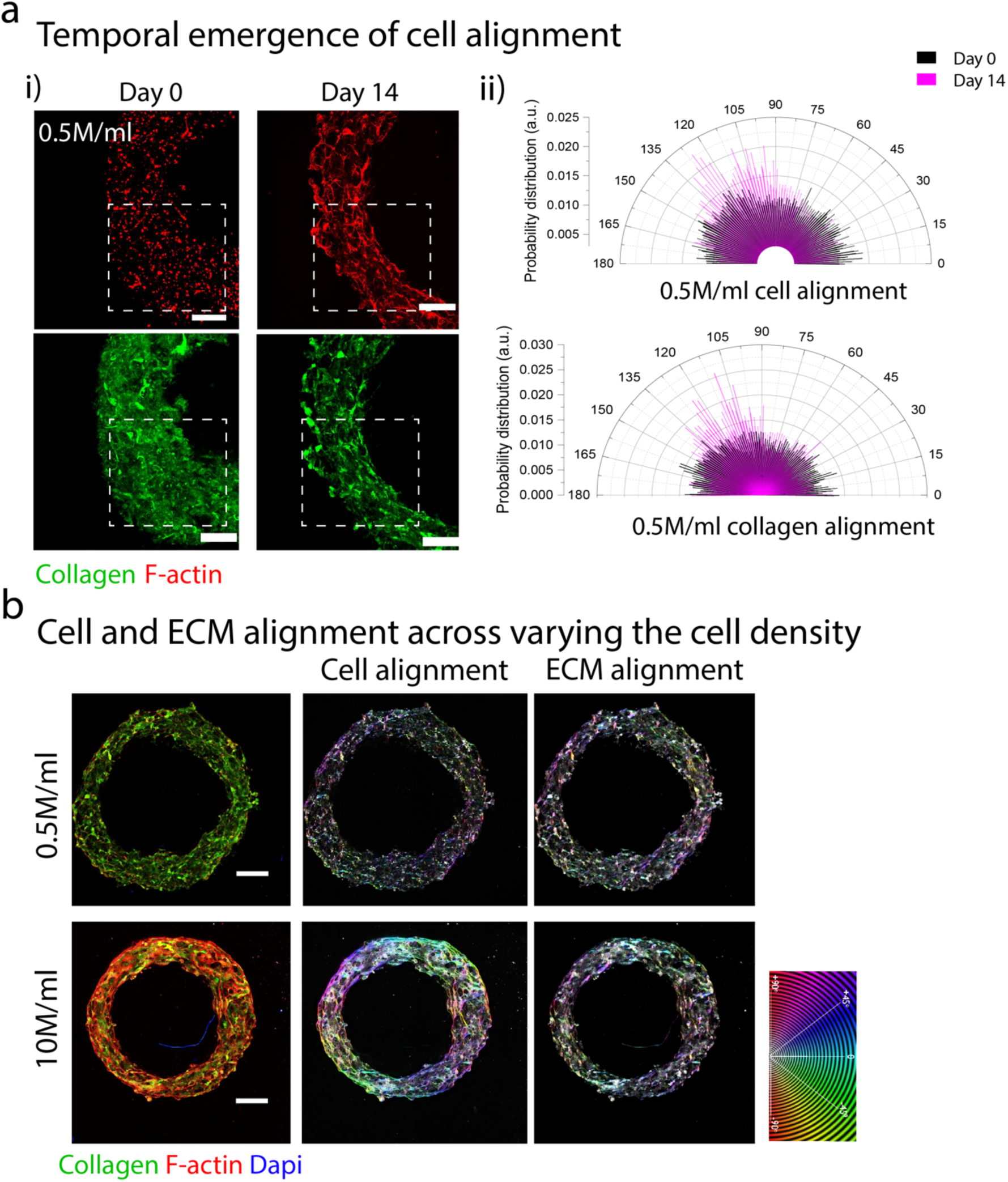
Influence of shape-morphing on cell and ECM alignment: **(a)** Temporal emergence of cell and ECM alignment in morphing tubes using 0.5M/ml cell density demonstrated by (i) confocal images of cells (top) and collagen fibres (bottom) on day 0 compared to day 14 (scale bar 500µm) and (ii) polar histogram showing the probable distribution of cells (top) and collagen fibres (bottom). **(b)** Confocal imaging of entire tubes for (i) 0.5M/ml (top), and (ii) 10M/ml (bottom) cell densities on day 14, and colour mapping images demonstrating dominant directions of cell and ECM alignment (Scale bar 500µm). All biological replicates n=3.

**Figure S7.**
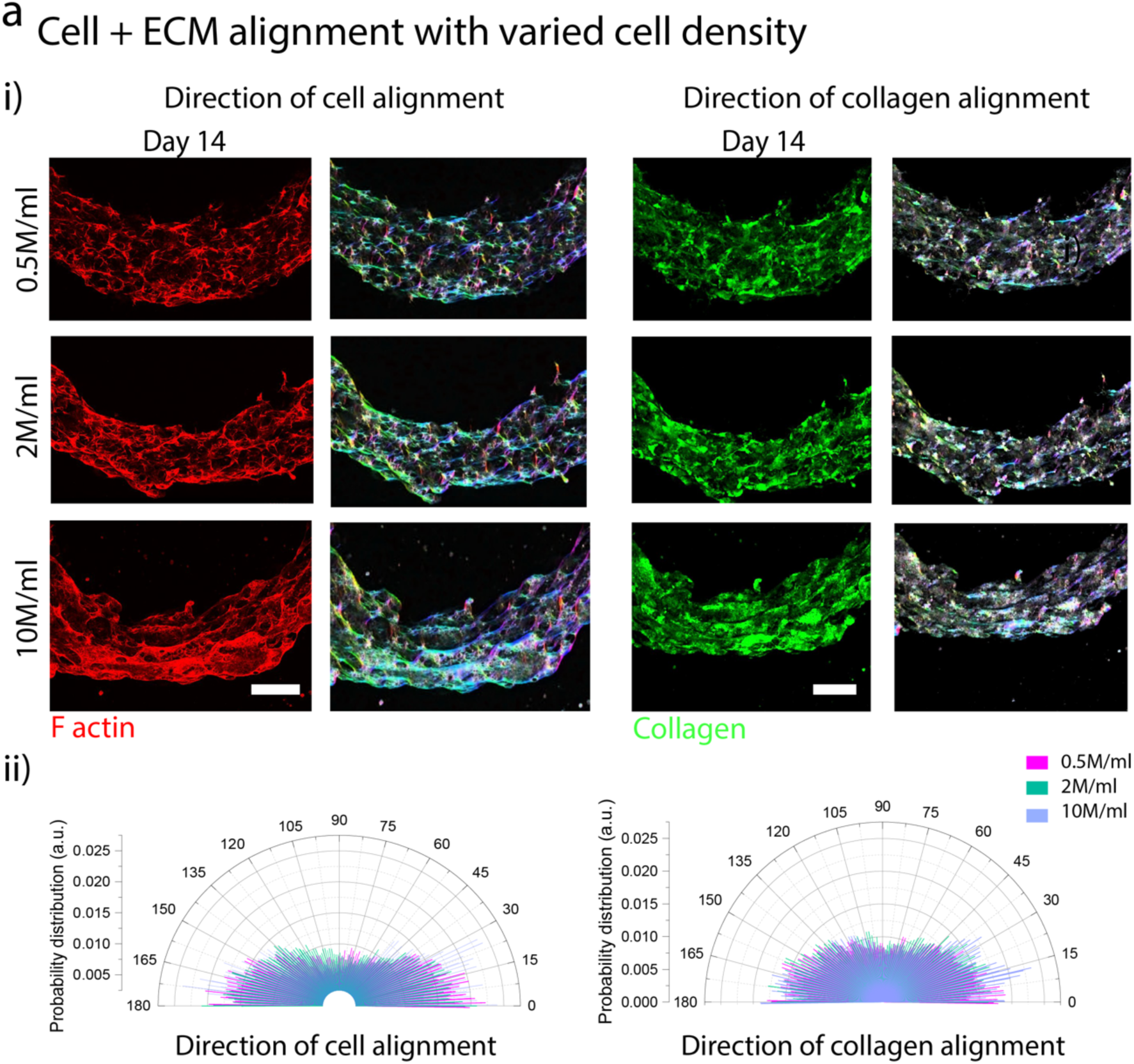
Cell and ECM alignment in shape-morphing tubes with varied cell density: **(a)** (i) Colour maps of cell (left) and ECM (right) alignment extracted from confocal images of F-actin and collagen within morphing tubes at varied cell densities (Scale bars 500µm). (ii) Quantitative assessment of cell and ECM alignment relative to the horizontal plane demonstrated through polar histograms (n=3 biological replicates)

**Figure S8.**
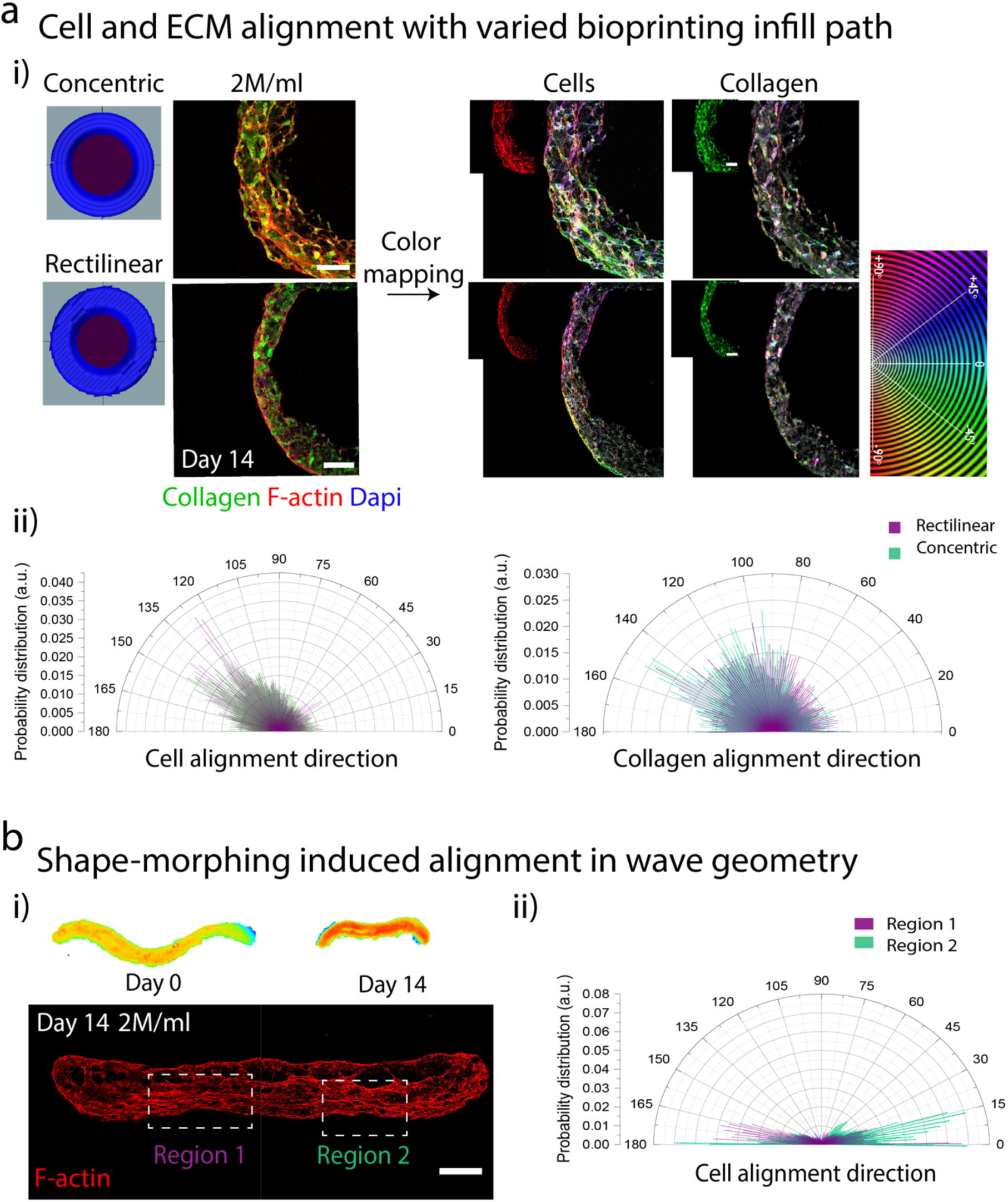
Cell alignment in morphing tubes with varied print path designs and geometry: **(a)** Cell and ECM alignment in shape-morphing tubes bioprinted using different print paths (concentric vs rectilinear infill) assessed by (i) confocal images of F-actin and collagen at day 14 followed by colour mapping to assess the dominant alignment directions. (ii) Polar histograms of cell and ECM alignment for tubes printed using concentric and rectilinear infills (n=3 biological replicates). **(b)** Cell alignment in shape-morphing wave geometry demonstrated by (i) Coloured brightfield images at day 0 and day 14 (scale bar 1mm) and confocal images of F-actin at day 14 (scale bar 500µm), and (ii) Cell alignment in regions 1 and 2 of the shape-morphing tissue plotted using polar histograms (n=3 biological replicates).

**Figure S9.**
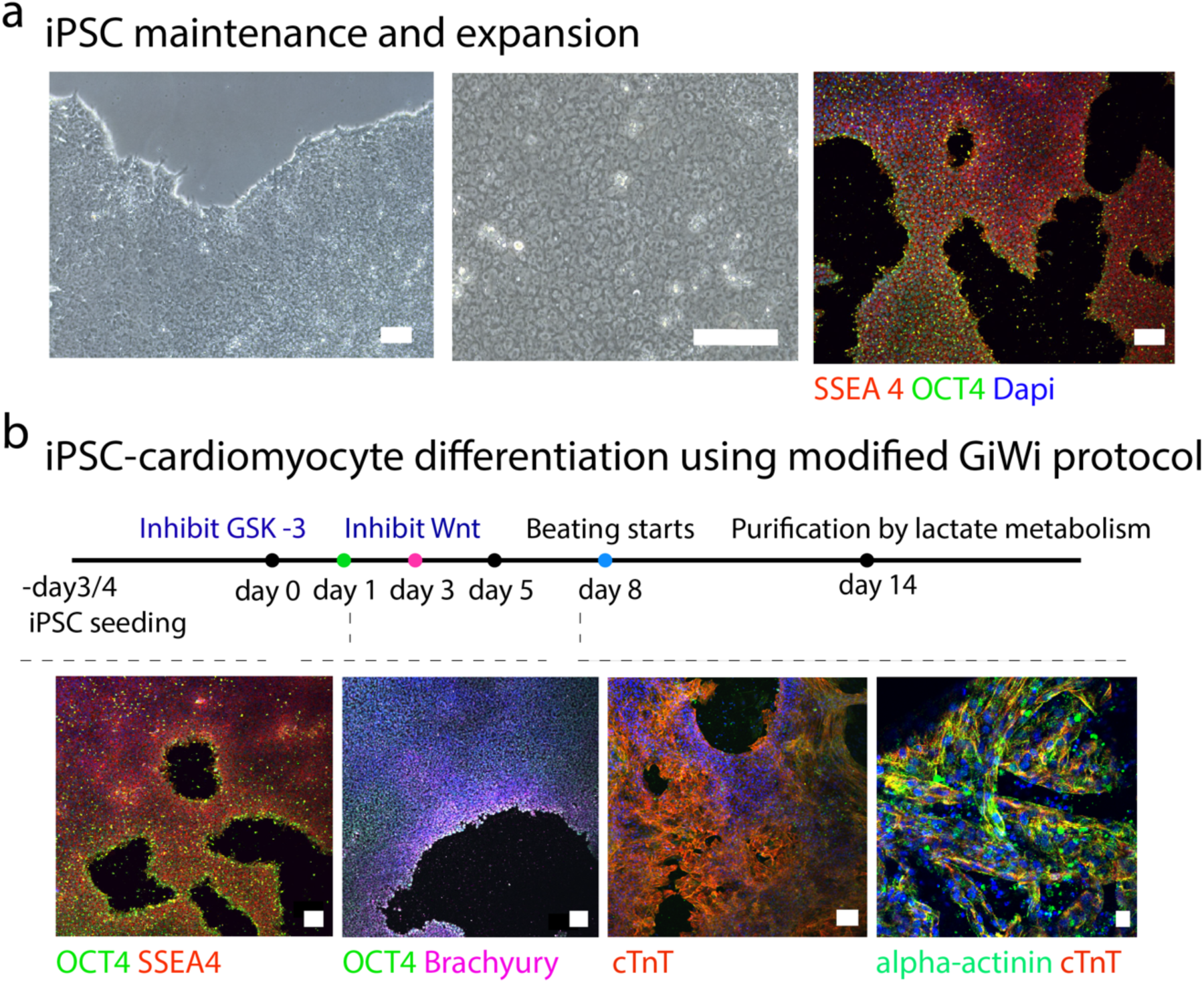
iPSC differentiation to cardiomyocytes: **(a)** Expansion and maintenance of iPSCs confirmed by (i) brightfield images and immunostaining for OCT4, SSEA4 (scale bar 100µm). **(b)** iPSC differentiated into cardiomyocytes using a modified GiWi protocol and purification based on lactate metabolism (OCT4/SSEA4, OCT4/Brachyury, cTnT scale bars all 100µm; alpha-actinin/cTnT scale bar 20µm).

**Figure S10.**
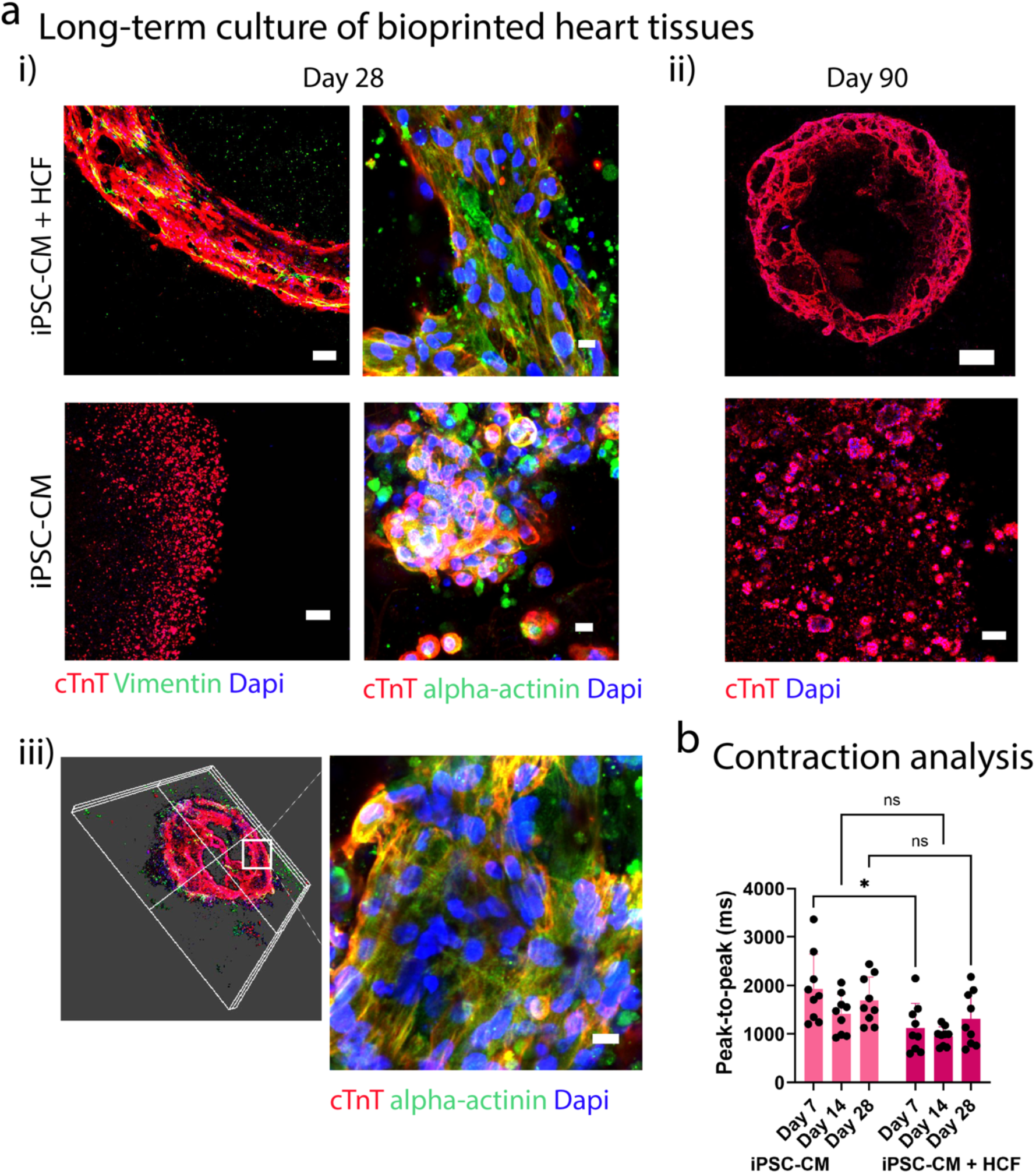
Bioprinting of shape-morphing iPSC-derived heart tissues: **(a)** Cell morphology within shape-morphing heart tubes (iPSC-CM + CF) compared to static controls (iPSC-CM only) assessed using confocal imaging on (i) day 28 and (ii) day 90 (Scale bars 100µm for cTnT/vimentin; 10µm for alpha-actinin/cTnT; 500µm for cTnT at day 90). (iii) 3D reconstruction of shape-morphing heart construct containing two chambers, and sarcomere staining within the heart wall (scale bar 10um). **(b)** Peak-to-peak time analysis of contraction curves for shape-morphing heart tubes (iPSC-CM + CF) compared to static controls (iPSC-CM only) at day 7,14 and 28 (n=9 biological replicates, Two-way ANOVA with Tukey’s multiple comparison test, where ns denotes not significant, and * denotes p < 0.05).

## Supporting Information S2 – Computational Framework

To investigate the active contractility and remodelling of contractile bioprinted tissue, we adapt the models of Reynolds et al.^32^ and Vigliotti et al. ^57^ to describe the thermodynamically consistent kinetics of stress fibre (SF) formation and the generation of active contractility. This framework considers key aspects of SF behaviour, including the thermodynamically-driven formation and dissociation of SFs, stress, strain and strain-rate induced remodelling, and conservation of cytoskeletal proteins within the cell. A representative volume element (RVE) in the undeformed state is defined as a sphere of radius *n^R^l*_0_/2. SFs emanate from the centre of this sphere, each comprised of *n^R^* in-series sarcomeres of length *l*_0_ in their ground state. SFs can form in a large number of initially uniformly distributed directions *n_θ_*. Sarcomeres have a steady state strain *ε̃_ss_* that minimizes their internal energy. When SFs experience a strain *ε_n,i_* in direction *i*, sarcomeres are added or removed until the steady state strain is achieved. The steady state number of sarcomeres in direction *i* is then

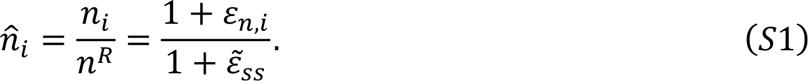

The steady state strain is determined from the real positive solution to

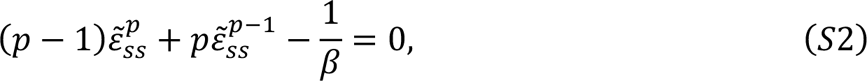

where *p* and *β* are constants that determine the internal energy of a sarcomere. Conservation of the total number of cytoskeletal proteins within each cell is enforced whereby

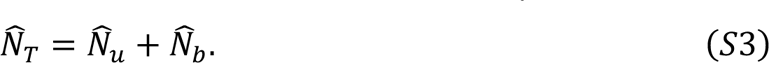

Where 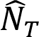 is the normalised total number of cytoskeletal proteins, 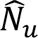, 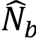 is the normalised number of unbound available proteins and bound within stress fibres respectively. The number of bound proteins can be determined as

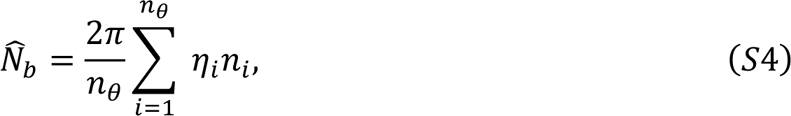

where *η̂_i_* is the steady state angular concentration of stress fibres, described by

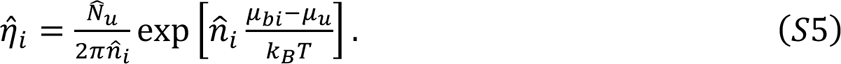

*μ_u_* is the standard enthalpy of the unbound proteins, *T* is the absolute temperature, *k_B_* is the Boltzmann constant, and *μ_bi_* is the standard enthalpy of a stress fibre with *n^R^* sarcomeres, given as:

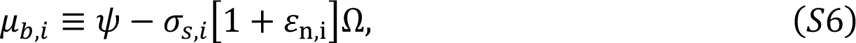

where Ω is the volume of a stress fibre with *n^R^* sarcomeres, and *Ψ* is the internal energy given as *Ψ* = *μ*_*b*0_ + *βμ*_*b*0_|*ε̃_ss_*|^*p*^. As we implement a steady state solution, the active stress within a SF is equal to the maximum isometric stress *σ_max_*. The active Cauchy stress tensor follows as:

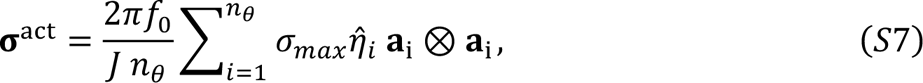

where *f*_0_ is the volume fraction of fibroblast cells and cytoskeletal proteins, *J* is the determinant of the deformation gradient **F**, and **a**_i_ = **Fa**_i,)_ where ***a***_i,0_ is a unit vector in direction *i*.

To describe the behaviour of initially randomly oriented collagen fibres in the bioprinted tissue, we implement a discrete fibre dispersion model adapted from **Holz and Odgen** ^58^ and developed by **McEvoy et al.** ^59^. Following from the cell model, we consider that fibres can exist in many discrete directions *m_θ_*. The passive collagen stress in a given direction *σ_f,j_* is ^60^

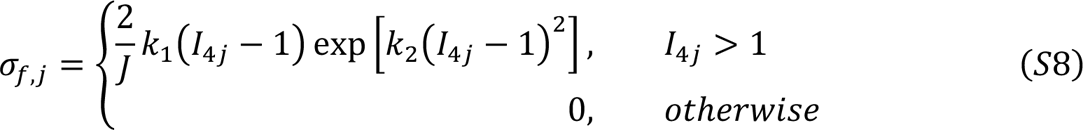

where *I*_4*j*_ is an invariant that represents the squared stretch in the direction of the unit vector ***a***_0*j*_ calculated as *I*_4*j*_ = **a**_j,0_ ⋅ (*Ca*_j,0_) with **C** the right Cauchy-Green tensor. *k*_1_ and *k*_2_ are material constants associated with fibre stiffening. In compression, the bulk behaviour of the tissue is captured by the underlying neo-Hookean model. The total Cauchy stress tensor for the dispersed collagen model follows as

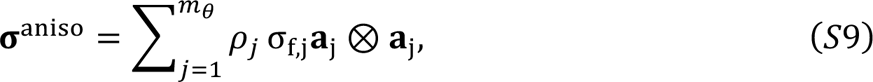

where *ρ_j_* is the fibre density in direction *i* and σ_f,j_ is the fibre bundle stress in that direction. The probability distribution for dispersed collagen is described by a von Mises distribution ^58^,

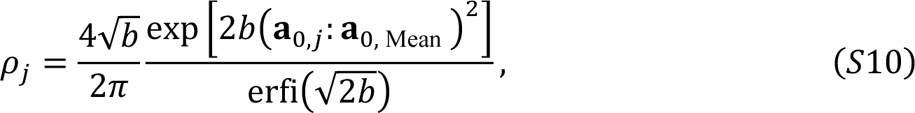

where *b* is a constant dispersion parameter, erfi(*x*) denotes the imaginary error function ***a***_0,*mean*_ is a unit vector indicating the mean fibre directions, and ***a***_0,*j*_ is a unit vector indicating one of *m_θ_* directions. The distribution is normalised such that 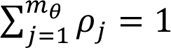. Here we choose the dispersion parameters such that there is initially a uniform distribution of collagen. The isotropic behaviour of the bioprinted tissue is described by a neo-Hookean hyperelastic model with a Cauchy stress given by:

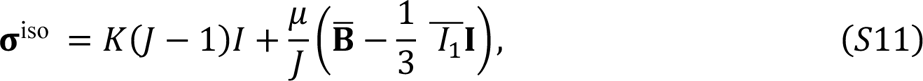

where ***B*** is the left Cauchy-Green tensor, and 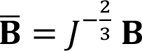 is the first invariant of **B** with *I*_1_ =tr(**B**) and 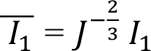. *μ* and *K* are the bulk and shear moduli. The mechanical behaviour of the granular hydrogel is also described by a neo-Hookean model. The material’s Young’s Modulus is determined by *E* = 9*Kμ*/(3*K* + *μ*).

Model parameters for the bioprinted tissue constructs and support bath are given as follows: Based on the experimental cell concentration, the volume fraction of cells and their cytoskeletal proteins is *f*_0_ = 6.685 × 10^−2^ unless stated otherwise. Simulations are reported for cells at experimental temperature T = 310K (∼37◦C). The remaining parameters for the SF framework are confined within ranges reported by Vigliotti et al. ^57^ and Reynolds et al. ^32^ with *Ω* = 7.9258 × 10^−8^ *μm;*^3^, *β* = 1.2, *p* = 2, *μ*_*b*0_ = 9*k_B_T* and *μ_u_* = 8*k_B_T*. The maximum isomeric stress is *σ_max_* = 240 kPa ^61^. The number of sampled directions *n_θ_* = *m_θ_* = 240 led to a converged solution. Parameters *k*_1_ and *k*_2_ for the collagen fibres material constants are given values 1.15 kPa and 0.3 respectively. The bulk and shear moduli are calibrated to *K* = 0.238 kPa and *μ* = 0.2173 kPa. The mechanical properties of the support bath are captured with a shear modulus *μ* = 6.76 × 10^−3^ kPa and bulk modulus *K* = 0.167 kPa, unless otherwise specified.

## Supporting Information S3 – Additional Tables

**Table S1:** Cell types.

| Cell type | Catalog no. | Company |
| --- | --- | --- |
| Human Cardiac Fibroblasts (HCF) | C-12375 | Promocell |
| Human Umbilical Vein Endothelial Cell (HuVEC) | C-12200 | Promocell |
| Gibco™ episomal hiPSC line | A18945 | Fisher Scientific |

**Table S2:**
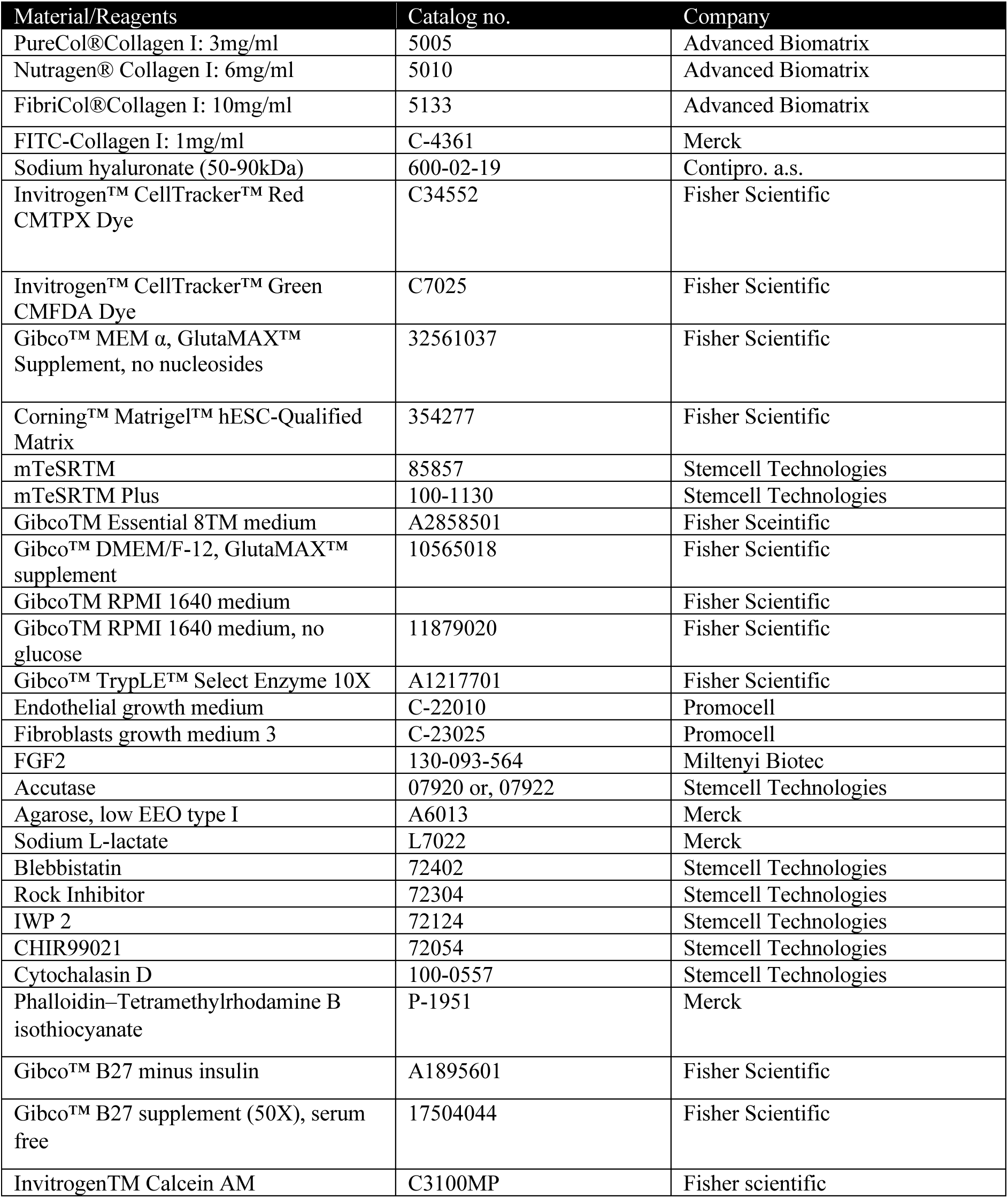
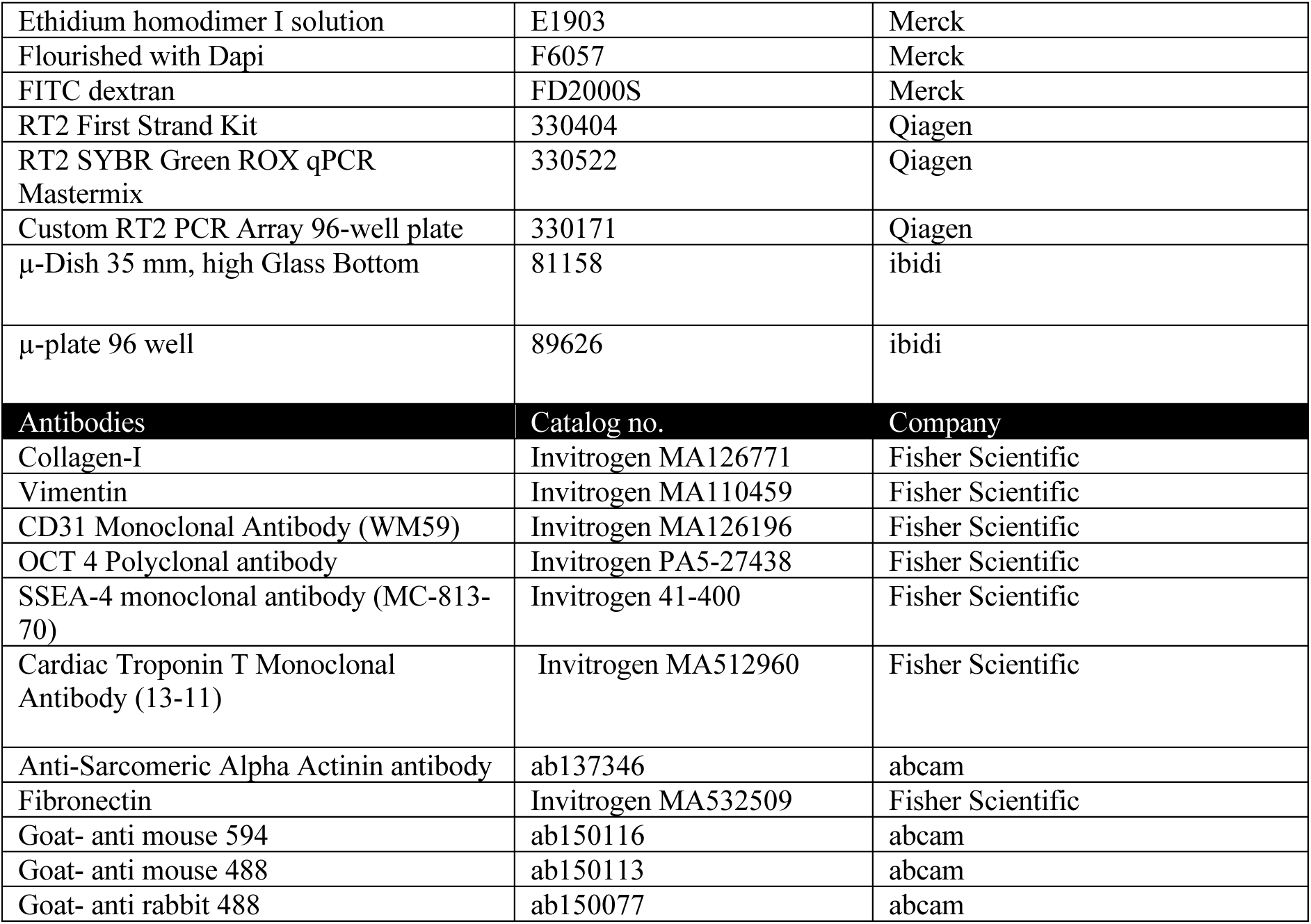
Materials and reagents.

**Table S3:**
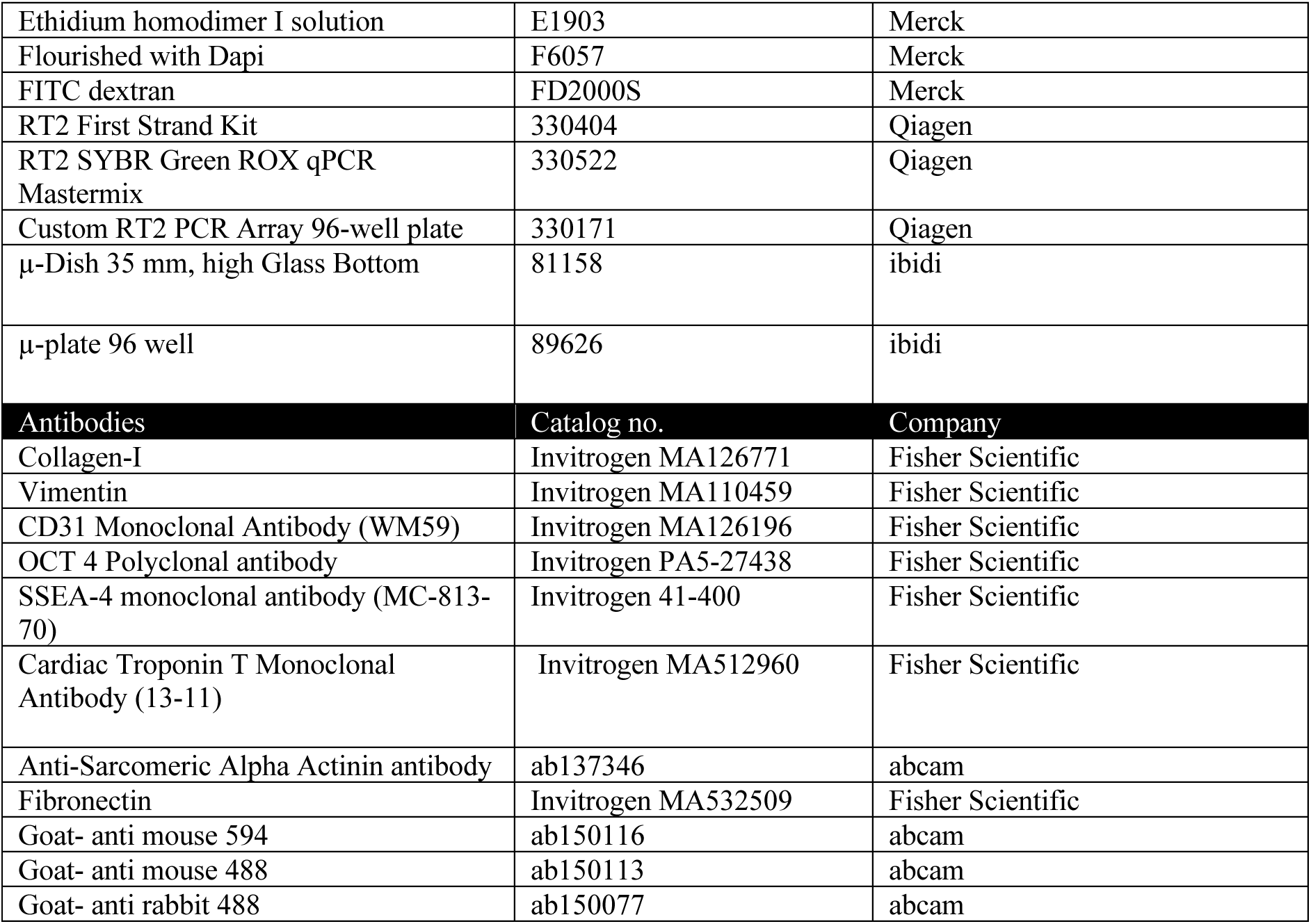
Custom RT2 PCR array gene details.

## References

1. Murphy, S. V & Atala, A. 3D bioprinting of tissues and organs. Nature Biotechnology 32, 773–785 (2014).

2. Skylar-Scott, M. A. et al. Biomanufacturing of organ-specific tissues with high cellular density and embedded vascular channels. Science Advances 5, eaaw2459 (2019).

3. Kolesky, D. B. et al. 3D Bioprinting of Vascularized, Heterogeneous Cell-Laden Tissue Constructs. Advanced Materials 26, 3124–3130 (2014).

4. Daly, A. C., Prendergast, M. E., Hughes, A. J. & Burdick, J. A. Bioprinting for the Biologist. Cell 184, 18–32 (2021).

5. Bagrat, G. et al. Multivascular networks and functional intravascular topologies within biocompatible hydrogels. Science 364, 458–464 (2019).

6. Sun, W. et al. The bioprinting roadmap. Biofabrication 12, 22002 (2020).

7. Bernal, P. N. et al. Volumetric Bioprinting of Complex Living-Tissue Constructs within Seconds. Advanced Materials 31, 1904209 (2019).

8. Ozbolat, I. T. & Hospodiuk, M. Current advances and future perspectives in extrusion-based bioprinting. Biomaterials 76, 321–343 (2016).

9. Lee, A. et al. 3D bioprinting of collagen to rebuild components of the human heart. Science 365, 482–487 (2019).

10. Hinton, T. J. et al. Three-dimensional printing of complex biological structures by freeform reversible embedding of suspended hydrogels. Science Advances 1, e1500758 (2015).

11. Esser, T. U. et al. Direct 3D-Bioprinting of hiPSC-Derived Cardiomyocytes to Generate Functional Cardiac Tissues. Advanced Materials 35, 2305911 (2023).

12. Malda, J. et al. 25th Anniversary Article: Engineering Hydrogels for Biofabrication. Advanced Materials 25, 5011–5028 (2013).

13. Levato, R. et al. From Shape to Function: The Next Step in Bioprinting. Advanced Materials 32, 1906423 (2020).

14. Stooke-Vaughan, G. A. & Campàs, O. Physical control of tissue morphogenesis across scales. Current Opinion in Genetics & Development 51, 111–119 (2018).

15. Kuang, X. et al. Advances in 4D Printing: Materials and Applications. Advanced Functional Materials 29, 1805290 (2019).

16. Ding, Z. et al. Direct 4D printing via active composite materials. Science Advances 3, e1602890 (2017).

17. Sydney Gladman, A., Matsumoto, E. A., Nuzzo, R. G., Mahadevan, L. & Lewis, J. A. Biomimetic 4D printing. Nature Materials 15, 413–418 (2016).

18. Momeni, F., M. Mehdi Hassani. N, S., Liu, X. & Ni, J. A review of 4D printing. Materials & Design 122, 42–79 (2017).

19. Gao, B. et al. 4D Bioprinting for Biomedical Applications. Trends in Biotechnology 34, 746–756 (2016).

20. Ashammakhi, N. et al. Advances and Future Perspectives in 4D Bioprinting. Biotechnology journal 13, e1800148–e1800148 (2018).

21. Wan, Z., Zhang, P., Liu, Y., Lv, L. & Zhou, Y. Four-dimensional bioprinting: Current developments and applications in bone tissue engineering. Acta Biomaterialia 101, 26–42 (2020).

22. Amukarimi, S. & Mozafari, M. 4D bioprinting of tissues and organs. Bioprinting 23, e00161 (2021).

23. Lee, Y. Bin et al. Induction of Four-Dimensional Spatiotemporal Geometric Transformations in High Cell Density Tissues via Shape-Changing Hydrogels. Advanced Functional Materials 31, 2010104 (2021).

24. Kirillova, A., Maxson, R., Stoychev, G., Gomillion, C. T. & Ionov, L. 4D Biofabrication Using Shape-Morphing Hydrogels. Advanced Materials 29, 1703443 (2017).

25. Gong, J. et al. Complexation-induced resolution enhancement of 3D-printed hydrogel constructs. Nature Communications 11, 1267 (2020).

26. Morley, C. D. et al. Quantitative characterization of 3D bioprinted structural elements under cell generated forces. Nat Commun 10, 3029 (2019).

27. Davidson, M. D. et al. Programmable and contractile materials through cell encapsulation in fibrous hydrogel assemblies. Science Advances 7, eabi8157 (2022).

28. Morley, C. D. et al. Quantitative characterization of 3D bioprinted structural elements under cell generated forces. Nature Communications 10, 1–9 (2019).

29. Polacheck, W. J. & Chen, C. S. Measuring cell-generated forces: a guide to the available tools. Nat Methods 13, 415–423 (2016).

30. Mailand, E., Li, B., Eyckmans, J., Bouklas, N. & Sakar, M. S. Surface and Bulk Stresses Drive Morphological Changes in Fibrous Microtissues. Biophysical Journal 117, 975–986 (2019).

31. Boudou, T. et al. A Microfabricated Platform to Measure and Manipulate the Mechanics of Engineered Cardiac Microtissues. Tissue Engineering Part A 18, 910–919 (2012).

32. Reynolds, N. H., McEvoy, E., Panadero Pérez, J. A., Coleman, R. J. & McGarry, J. P. Influence of multi-axial dynamic constraint on cell alignment and contractility in engineered tissues. Journal of the Mechanical Behavior of Biomedical Materials 112, 104024 (2020).

33. Heer, N. C. & Martin, A. C. Tension, contraction and tissue morphogenesis. Development 144, 4249–4260 (2017).

34. Prendergast, M. E., Davidson, M. D. & Burdick, J. A. A Biofabrication Method to Align Cells within Bioprinted Photocrosslinkable and Cell-degradable Hydrogel Constructs via Embedded Fibers. Biofabrication 13, 10.1088/1758-5090/ac25cc (2021).

35. Kim, H. et al. Shear-induced alignment of collagen fibrils using 3D cell printing for corneal stroma tissue engineering. Biofabrication 11, 035017 (2019).

36. Ahrens, J. H. et al. Programming Cellular Alignment in Engineered Cardiac Tissue via Bioprinting Anisotropic Organ Building Blocks. Advanced Materials 34, 2200217 (2022).

37. Veerman, C. C. et al. Immaturity of Human Stem-Cell-Derived Cardiomyocytes in Culture: Fatal Flaw or Soluble Problem? Stem Cells and Development 24, 1035–1052 (2015).

38. Le Garrec, J. F. et al. A predictive model of asymmetric morphogenesis from 3D reconstructions of mouse heart looping dynamics. eLife 6, 1–35 (2017).

39. Kawahira, N., Ohtsuka, D., Kida, N., Hironaka, K. ichi & Morishita, Y. Quantitative Analysis of 3D Tissue Deformation Reveals Key Cellular Mechanism Associated with Initial Heart Looping. Cell Reports 30, 3889–3903.e5 (2020).

40. Mandrycky, C. J. et al. Engineering Heart Morphogenesis. Trends in Biotechnology 38, 835–845 (2020).

41. Zhang, P., Su, J. & Mende, U. Cross talk between cardiac myocytes and fibroblasts: from multiscale investigative approaches to mechanisms and functional consequences. Am J Physiol Heart Circ Physiol 303, H1385–H1396 (2012).

42. Zhou, P. & Pu, W. T. Recounting cardiac cellular composition. Circ Res 118, 368–370 (2016).

43. Goldfracht, I. et al. Generating ring-shaped engineered heart tissues from ventricular and atrial human pluripotent stem cell-derived cardiomyocytes. Nat Commun 11, 75 (2020).

44. Sala, L. et al. MUSCLEMOTION: A Versatile Open Software Tool to Quantify Cardiomyocyte and Cardiac Muscle Contraction In Vitro and In Vivo. Circ Res 122, e5– e16 (2018).

45. Giacomelli, E. et al. Human-iPSC-Derived Cardiac Stromal Cells Enhance Maturation in 3D Cardiac Microtissues and Reveal Non-cardiomyocyte Contributions to Heart Disease. Cell Stem Cell 26, 862–879.e11 (2020).

46. Beauchamp, P. et al. 3D Co-culture of hiPSC-Derived Cardiomyocytes With Cardiac Fibroblasts Improves Tissue-Like Features of Cardiac Spheroids. Front. Mol. Biosci. 7, (2020).

47. Hookway, T. A. et al. Bi-directional Impacts of Heterotypic Interactions in Engineered 3D Human Cardiac Microtissues Revealed by Single-Cell RNA-Sequencing and Functional Analysis. bioRxiv 2020.07.06.190504 (2020) doi:10.1101/2020.07.06.190504.

48. Ronaldson-Bouchard, K. et al. Advanced maturation of human cardiac tissue grown from pluripotent stem cells. Nature 556, 239–243 (2018).

49. Lian, X. et al. Directed cardiomyocyte differentiation from human pluripotent stem cells by modulating Wnt/β-catenin signaling under fully defined conditions. Nat Protoc 8, 162– 175 (2013).

50. Zhao, M., Tang, Y., Zhou, Y. & Zhang, J. Deciphering Role of Wnt Signalling in Cardiac Mesoderm and Cardiomyocyte Differentiation from Human iPSCs: Four-dimensional control of Wnt pathway for hiPSC-CMs differentiation. Sci Rep 9, 19389 (2019).

51. Horikoshi, Y. et al. Fatty Acid-Treated Induced Pluripotent Stem Cell-Derived Human Cardiomyocytes Exhibit Adult Cardiomyocyte-Like Energy Metabolism Phenotypes. Cells 8, 1095 (2019).

52. Voges, H. K. et al. Development of a human cardiac organoid injury model reveals innate regenerative potential. Development dev.143966 (2017) doi:10.1242/dev.143966.

53. Senior, J. J., Cooke, M. E., Grover, L. M. & Smith, A. M. Fabrication of Complex Hydrogel Structures Using Suspended Layer Additive Manufacturing (SLAM). Advanced Functional Materials 29, 1904845 (2019).

54. Mirdamadi, E., Muselimyan, N., Koti, P., Asfour, H. & Sarvazyan, N. Agarose Slurry as a Support Medium for Bioprinting and Culturing Freestanding Cell-Laden Hydrogel Constructs. 3D Printing and Additive Manufacturing 6, 158–164 (2019).

55. D. Clemons, T., et al. Coherency image analysis to quantify collagen architecture: implications in scar assessment. RSC Advances 8, 9661–9669 (2018).

56. Sala, L. et al. Musclemotion. Circulation Research 122, e5–e16 (2018).

57. Vigliotti, A., Ronan, W., Baaijens, F. P. T. & Deshpande, V. S. A thermodynamically motivated model for stress-fiber reorganization. Biomech Model Mechanobiol 15, 761– 789 (2016).

58. Holzapfel, G. A. & Ogden, R. W. On the tension–compression switch in soft fibrous solids. European Journal of Mechanics - A/Solids 49, 561–569 (2015).

59. McEvoy, E., Holzapfel, G. A. & McGarry, P. Compressibility and Anisotropy of the Ventricular Myocardium: Experimental Analysis and Microstructural Modeling. Journal of Biomechanical Engineering 140, (2018).

60. Nolan, D. R., Gower, A. L., Destrade, M., Ogden, R. W. & McGarry, J. P. A robust anisotropic hyperelastic formulation for the modelling of soft tissue. Journal of the Mechanical Behavior of Biomedical Materials 39, 48–60 (2014).

61. Lucas, S. M., Ruff, R. L. & Binder, M. D. Specific tension measurements in single soleus and medial gastrocnemius muscle fibers of the cat. Experimental Neurology 95, 142–154 (1987).

